# Targeted, Genome-scale Overexpression in Proteobacteria

**DOI:** 10.1101/2024.03.01.582922

**Authors:** Amy B. Banta, Ashley N. Hall, Kevin S. Myers, Ryan D. Ward, Bryce C. Davis, Rodrigo A. Cuellar, Michael Place, Claire C. Freeh, Emily E. Bacon, Jason M. Peters

## Abstract

Targeted, genome-scale gene perturbation screens using Clustered Regularly Interspaced Short Palindromic Repeats interference (CRISPRi) and activation (CRISPRa) have revolutionized eukaryotic genetics, advancing medical, industrial, and basic research. Although CRISPRi knockdowns have been broadly applied in bacteria, options for genome-scale gene overexpression face key limitations. Here, we develop a facile approach for genome-scale overexpression in bacteria we call, “CRISPRtOE” (CRISPR transposition and OverExpression). We first create a platform for comprehensive gene targeting using CRISPR-associated transposons (CAST) and show that transposition occurs at a higher frequency in non-transcribed DNA. We then demonstrate that CRISPRtOE can upregulate gene expression in Proteobacteria with medical and industrial relevance by integrating synthetic promoters of varying strength upstream of target genes. Finally, we employ CRISPRtOE screening at the genome-scale in the model bacterium *Escherichia coli* and the non-model biofuel producer *Zymomonas mobilis*, recovering known and novel antibiotic and engineering targets. We envision that CRISPRtOE will be a valuable overexpression tool for antibiotic mode of action, industrial strain optimization, and gene function discovery in bacteria.

**Importance:** Systematic alteration of bacterial gene expression enables identification of genes relevant to diverse fields of study and practical applications. Although many targeted, genome-scale genetic tools exist for reducing or eliminating gene expression, there are few facile approaches for systematic gene overexpression in bacteria. Here, we develop a targeted overexpression approach for Proteobacteria of medical and industrial importance that precisely inserts strong promoters upstream of genes using CRISPR-associated transposons. We demonstrate that this approach can be used to systematically overexpress genes in both model (*Escherichia coli* K-12) and non-model (*Zymomonas mobilis*) Proteobacteria for the purpose of understanding antibiotic resistance mechanisms and improving strain resilience in biofuel production conditions, respectively.

## Introduction

Overexpression (OE) studies have been invaluable for gene phenotyping in medical (1–3), industrial (4), and basic contexts (3, 5). When used at the genome-scale, OE enables systematic functional interrogation of genes that are normally cryptic under screening conditions, mimics natural resistance mechanisms, and complements loss of function approaches by identifying genes with opposite phenotypes when knocked down versus overexpressed (1, 6). Clustered Regularly Interspaced Short Palindromic Repeats activation (CRISPRa) has allowed routine, genome-scale overexpression screening in eukaryotes for drug synergy/antagonism (6) or sensitivity/resistance to toxic plant compounds that reduce bioproduct yields (7). For example, a recent screen in mammalian cells for genes that interact with the anti-cancer drug, rigosertib, leveraged genome-scale CRISPR interference (CRISPRi) knockdowns with CRISPRa OE to identify target pathways (1). OE has also enhanced strain optimization for industrial purposes, such as a genome-wide CRISPRa screen in in *Saccharomyces cerevisiae* for genes involved in resistance to the biofuel production stressor, furfural (7).

Existing genome-scale OE approaches in bacteria have key limitations. Classic OE approaches, such as random integration of promoter-containing transposons to overexpress downstream genes(8), are untargeted and require high-density transposon (Tn) libraries to resolve phenotypic ambiguities caused by the effects of Tn insertion versus OE. Libraries of Open Reading Frames (ORFs) cloned into multi-copy plasmids (9) are technically targeted approaches, but are tedious and expensive to construct, limiting their utility in non-model bacteria. Further, the use of plasmids presents additional complications, such as the constant requirement for antibiotic selection for plasmid maintenance—complicating screens for antibiotic function—and copy number variation that increases experimental noise (10). Contemporary approaches have sought to mitigate these issues and make genome-scale OE more accessible. For instance, Trackable Multiplex Recombineering (TRMR) is a targeted approach that uses recombinase-aided DNA editing (recombineering) to insert barcoded promoters or ribosome binding variants upstream of genes to achieve OE (11, 12); however, use of this method at the genome-scale is restricted to bacteria with efficient recombineering systems. An innovative new OE technique, “Dub-seq,” uses random genomic fragments cloned into barcoded, multi-copy plasmids (13, 14). Dub-seq reduces upfront barriers to OE library construction, but at the cost of being untargeted, increasing the complication of downstream analysis, and sharing some downsides of other plasmid-based approaches. Moreover, the length of cloned genomic fragments dictates Dub-seq library screening results; smaller fragments may miss genes that only have phenotypes when co-expressed in operons (e.g., protein complex subunits) while longer fragments may make it more difficult to pinpoint individual genes responsible for the selected phenotype.

Although systematic CRISPRa screens are routinely used in eukaryotes (1–3), no genome-scale CRISPRa studies have been reported in bacteria, to our knowledge. CRISPRa in eukaryotes uses a catalytically inactive Cas9 protein (dCas9) and single guide RNAs (sgRNAs) to deliver activator proteins to promoter regions, facilitating activation by opening chromatin (15). In contrast, CRISPRa-based OE in bacteria requires direct interactions between activator proteins and RNA polymerase (RNAP) that dictate strict activator-RNAP spacing and orientation on DNA(16–19). Recent improvements to bacterial CRISPRa using dCas9 variants with reduced PAM stringency or engineered activator proteins (19) partially mitigated the spacing requirements and increased overall activation; however—even in these optimized systems— only 1/3 to 1/2 of targeted endogenous genes could be upregulated, and the extent of OE was unpredictable (18, 19).

Naturally occurring CRISPR-associated transposon (CAST) systems(20) have been engineered to generate targeted insertions in bacterial genomes with high efficiency(21–23). Although several CAST systems have been described (24–26), the Tn*6677* system from *Vibrio cholerae* (VcCAST) is among the best characterized and most precise(27–30). In VcCAST, a nuclease-deficient Type I-F CRISPR system pairs with a Tn*7*-like transposase to achieve guide RNA-directed transposition. Guide RNA spacers direct VcCAST proteins to a complementary protospacer, resulting in transposition of Tn*6677* end-flanked DNA ∼49 bp downstream of the protospacer (27–30). Transposon DNA can be inserted in two orientations with respect to the Tn*6677* ends (R-L or L-R) but is highly biased toward the R-L orientation (21, 31). VcCAST is highly precise, with nearly all tested guides showing >95% on-target insertion activity. The utility of CASTs has been demonstrated for high-efficiency genome editing in Proteobacteria (21), systematic disruption of transcription factor genes in *Pseudomonas aeruginosa* (23), and targeting individual strains in a complex microbiome (22), but existing systems remain sub-optimal for genome-scale functional genomics. Moreover, the use of Tn*6677* to deliver regulatory elements—such as promoters—has recently shown promise in the model bacterium, *Escherichia coli* (32). However, key gaps remain in our ability to deploy CAST systems for genome-scale OE studies in diverse bacteria.

Here we addressed those gaps by creating a VcCAST system for targeted, genome-scale OE in Proteobacteria we call, “CRISPRtOE” (CRISPR transposition and OverExpression) (Fig. 1). CRISPRtOE precisely delivers a transposon with an outward facing promoter upstream of genes to facilitate OE. We show the utility of CRISPRtOE to upregulate gene expression in medically and industrially-relevant Proteobacterial species. Finally, we demonstrate genome-scale overexpression in model (*E. coli*) and non-model (*Zymomonas mobilis*) Proteobacteria, defining genes required for resistance to antibiotics or biofuel production stressors, respectively.

**Figure 1.**
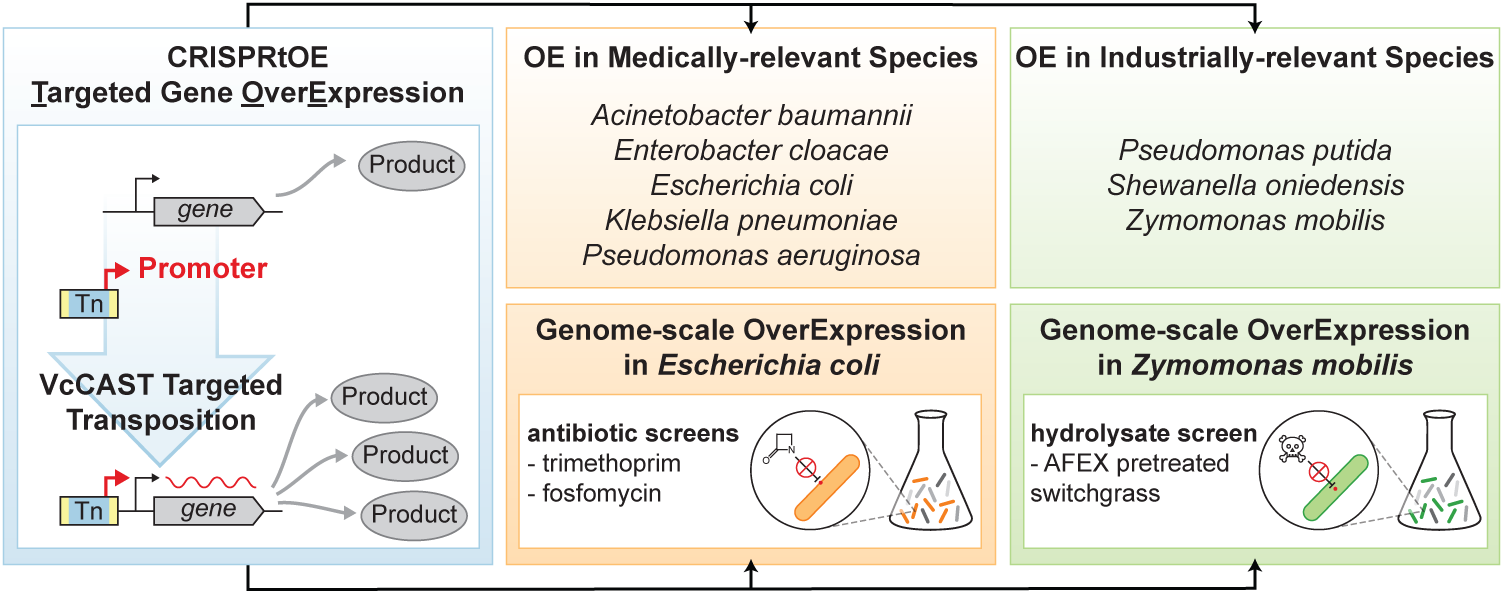
A CRISPR-associated transposition (CAST) system for targeted, genome-scale overexpression in Proteobacteria called “CRISPRtOE” (CRISPR transposition and OverExpression). CRISPRtOE precisely delivers a transposon with an outward facing promoter upstream of genes to facilitate overexpression. CRISPRtOE can upregulate gene expression in medically and industrially-relevant Proteobacterial species. Genome-scale CRISPRtOE screens in model (*E. coli*) and non-model (*Zymomonas mobilis*) Proteobacteria can be used to define genes required for resistance to antibiotics or biofuel production stressors.

## Results

### A VcCAST Platform for Genome-scale Functional Genomics

We first generated a VcCAST system designed for genome-scale functional genomics we call, “CRISPRt.” Like previous VcCAST constructs (21, 22), CRISPRt enables insertion of Tn*6677* at precise genomic positions specified by programmable gRNAs (Fig. 2A and 2B). Superimposed on this are additional properties for facile use of VcCAST at the genome-scale: 1) Tn insertion is selectable such that all transconjugants contain an integrated Tn; 2) *cas* helper and Tn plasmids only replicate in *pir*-containing *E. coli* cells and are non-replicative in recipient bacteria, obviating the need for plasmid curing prior to phenotyping (23); 3) *cas* and Tn components are on separate plasmids, preventing insertion of *cas* genes into the genome in cases of Tn plasmid co-integration and reducing Tn plasmid size which facilitates conjugation and cargo modification by standard cloning procedures (Fig. 2A, 2B, S1A, and S1B). CRISPRt transconjugants obtained from tri-parental matings were recovered at efficiencies consistent with genome-scale library creation (e.g., ∼10^-2^ frequency for *E. coli* K-12 MG1655, Fig. S1C). To confirm the precision of CRISPRt, we targeted both endogenous (*lacZ*) and heterologous (*gfp*) reporter genes for disruption in *E. coli* K-12 (Fig. S1D), using phenotypic assays and Illumina short read sequencing of the genomic DNA-Tn*6677* junction (Tn-seq (33)) as readouts (Fig. 2C, S2A, S2B, S2C, S2D, and S2E). As expected, Tn-seq showed a >99.5% on target efficiency for both *lacZ* and *gfp* spacers, Tn insertion within a tight window centered at ∼49 downstream of the protospacer, and a strong Tn insertion bias toward the R-L orientation. Tn plasmid co-integration occurred at a very low frequency (∼10^-3^ for *E. coli* K-12, Fig. S2F).

**Figure 2.**
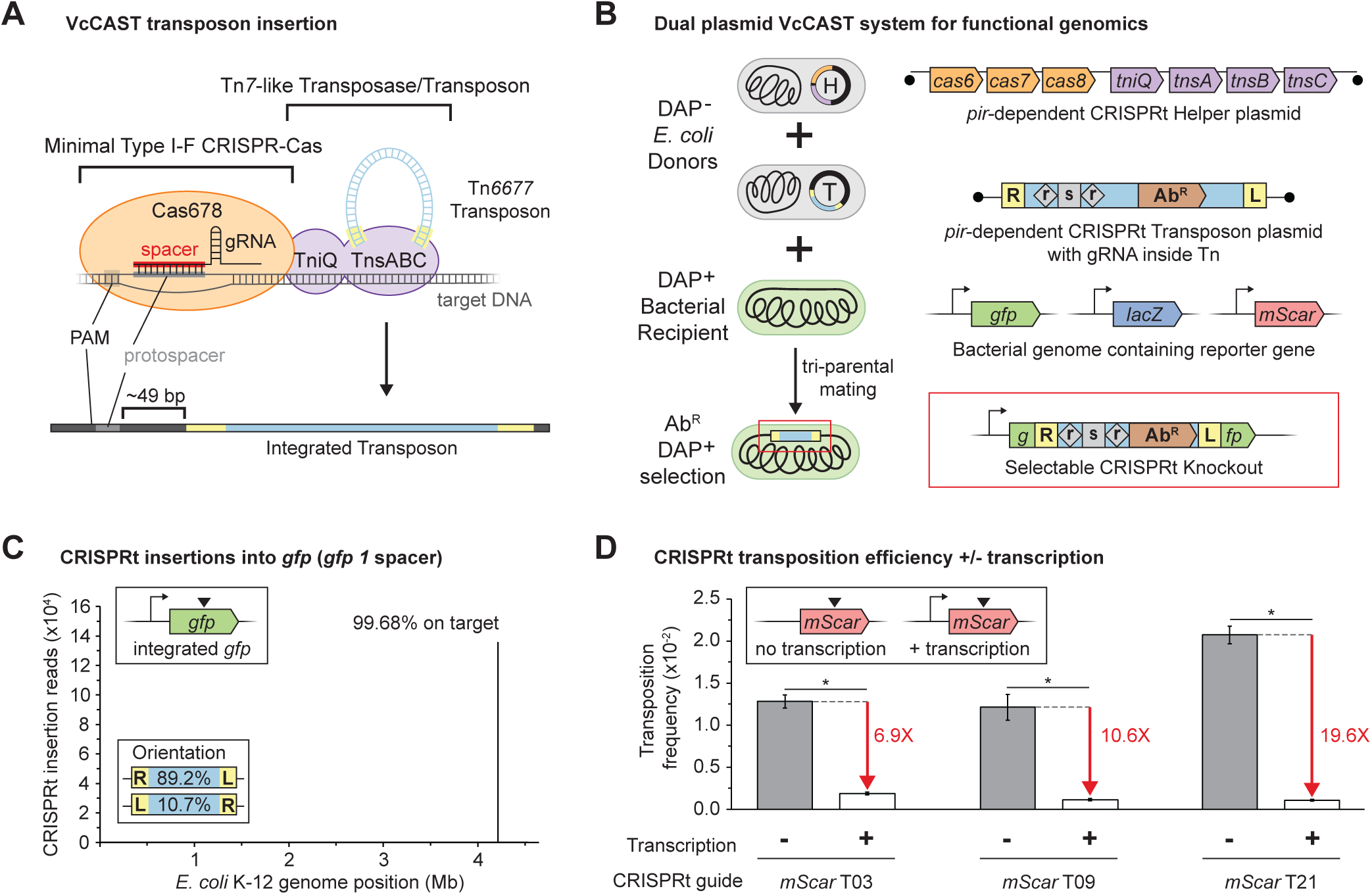
A mobilizable, selectable, dual-plasmid system for CRISPR-guided transposition. (A) Schematic of mechanism of *Vibrio cholera* Type I-F CRISPR-associated transposon (VcCAST) system(20, 21). (B) Schematic of strain construction using the CRISPRt targeted transposition system. Plasmids have pir-dependent origins of replication preventing replication in the bacterial recipient. One plasmid encodes the minimal Type I-F CRISPR-Cas (Cas678) and Tn*7*-like transposase (TnsABC and TniQ) machinery, and a second plasmid contains a transposon carrying the guide RNA and antibiotic resistance expression cassettes (see Figure S1 for details). These plasmids are transferred by co-conjugation to a recipient bacterium by *E. coli* donor strains with a chromosomal copy of the RP4 transfer machinery. Inside the recipient cell, the transposon (flanked in yellow) is inserted onto the chromosome at a site determined by the sequence of the guide RNA. Selection on antibiotic plates lacking diaminopimelic acid (DAP) selects for transconjugants and against the *E. coli* donor strains. (C) Specificity of CRISPRt disruption of *gfp* in *E. coli* measured by Tnseq. Mapping of location of transposon insertion sites to the *E. coli* genome after CRISPRt targeted transposition with the *gfp1* guide (226,671 reads). (D) Transposition efficiency of 3 individual CRISPRt guides (T03, T09, T21) targeting *mScarlet-I* cassettes in the *E. coli att*_Tn*7*_ site either with a promoter (+ transcription) or without a promoter (- transcription) measured by plating efficiency with and without selection (+/- kanamycin) in triplicate. Red arrows indicate fold change in efficiency without and with transcription. Error represents the average of n=3 assays, * indicates p<0.05, two-tailed t-test.

Although Protospacer Adjacent Motif (PAM) requirements for VcCAST have been well characterized (a “CN” PAM is sufficient (25)), other genomic features that impact guide efficacy are poorly understood. To address this issue, we created a pooled CRISPRt library targeting both strands of an *mScarlet-I* reporter gene integrated into the *E. coli* K-12 genome (46 gRNAs total, Fig. S3A). Surprisingly, we found that transcription had a significant impact on guide efficacy. To measure relative guide efficacy, we compared the abundance of gRNA spacers in our library before and after transposition into *mScarlet-I* (Fig. S3B). Consistent with previous data from smaller sets of spacers(34), we found considerable variability, with a ∼27-fold difference in efficacy between the most and least active guides. Looking for patterns that could explain these differences, we noticed a clear, but not absolute bias toward higher efficiency in template strand targeting guides. Other variables, such as distance from the gene start or PAM identity did not show an obvious trend (Fig. S3C and S3D). The strand bias in insertion frequency suggested either replication or transcription could be impacting guide efficacy. We continued to observe a strand bias when the orientation of *mScarlet-I* was flipped relative to the origin of replication (Fig. S4A and S4B), seemingly ruling out replication as a factor and implicating transcription. Indeed, CRISPRt insertion into a non-transcribed *mScarlet-I* gene showed reduced variation in relative guide efficacy (Fig. S5A). To determine the absolute effect of transcription on CRISPRt efficacy, we cloned the three most active guides from our *mScarlet-I* library (T03, T09, and T21) and individually tested their efficiencies in the presence and absence of transcription (Fig. 2E and S5B). We found that transcription of *mScarlet-I* reduced recovery of CRISPRt transconjugants by 6.9 to 19.6-fold, depending on the guide tested. We conclude that CRISPRt has excellent properties for genome-scale functional genomics while retaining the precision of VcCAST and that CRISPRt efficacy is higher when targeting non-transcribed DNA.

### Targeted and Tunable Overexpression in Proteobacteria

We sought to modify CRISPRt into a targeted overexpression system by inserting outward-facing promoters into Tn*6677* that increase transcription of downstream genes (CRISPRtOE (Fig. 3A)). We inserted synthetic, constitutive promoters of varying strength that had been shown to function in Gamma- and Alpha-proteobacteria (35, 36) into Tn*6677* with the goal of creating an OE gradient (Fig. S6A and S6B). To quantify the OE activity of CRISPRtOE, we generated a “test” strain with a promoterless *mScarlet-I* reporter gene integrated into the *E. coli* K-12 genome. This cassette contained a well-characterized protospacer upstream of *mScarlet-I* to act as a “landing pad” (LP) for CRISPRtOE transposons (Fig. 3A and S6C). The test strain showed negligible mScarlet-I expression in the absence of an upstream CRISPRtOE insertion (Fig. 3B).

**Figure 3.**
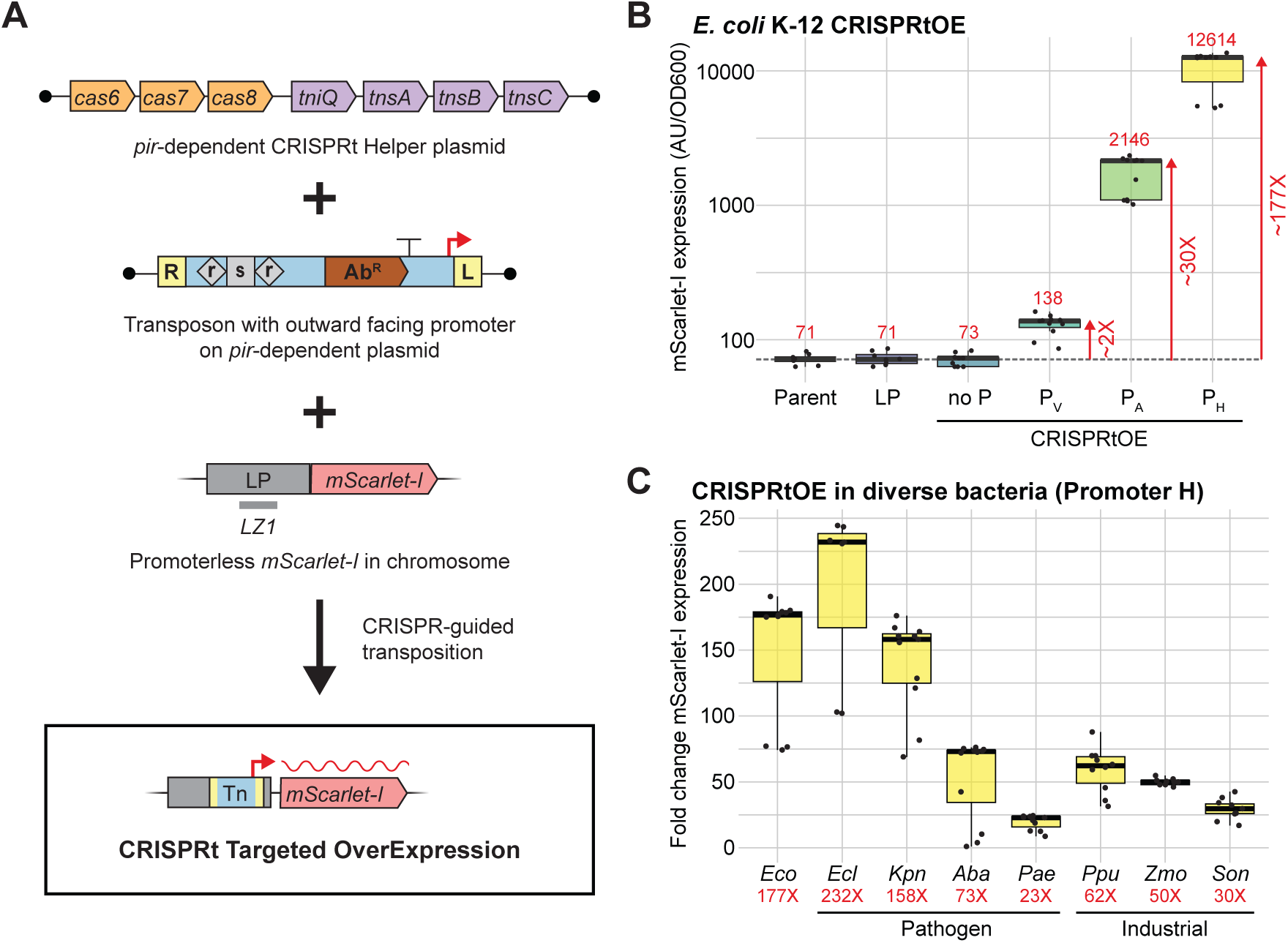
Tunable overexpression of chromosomally located genes using CRISPRtOE. (A) Schematic of the CRISPRtOE targeted overexpression system. The test chromosomal target is an *mScarlet-I* gene preceded by a ’Landing Pad’ (LP) with no promoter, a PAM, and the *LZ1* protospacer (see Figure S6C). The CRISPRtOE construct has a transposon carrying guide RNA and antibiotic resistance expression cassettes and an outward facing promoter (see Figure S6B for details). Co-conjugation of strains carrying the CRISPRtOE construct and the CRISPRt-H plasmid (harboring the VcCAST machinery) with a promoterless ’LP’ *mScarlet-I* reporter recipient strain results in a CRISPRtOE test strain. (B) mScarlet-I fluorescence analysis of *E. coli* CRISPRtOE isolates (no promoter or synthetic promoters A, V, and H) compared to the parent (promoterless ’LP’ *mScarlet-I*) strain. CRISPRtOE transposon insertion position and fluorescence measurements were determined for twelve isolates. Fluorescence measurements were normalized to cell density (OD_600_). On target, RL orientation isolates are shown here, see Figure S7B for all orientations and Table S6 for values. Error is expressed for the median value of 8-12 isolates in n=3 assays. (C) Fold effect of CRISPRtOE mScarlet-I overexpression using synthetic promoter H in eight Alpha- and Gammaprotebacteria (*E. coli* (*Eco*), *Enterobacter cloacae* (*Ecl*), *Klebsiella pneumoniae* (*Kpn*), *Acinetobacter baumannii* (*Aba*), *Pseudomonas aeruginosa* (*Pae*), *Pseudomonas putida* (*Ppu*), *Zymomonas mobilis* (*Zmo*), and *Shewanella oneidensis* (*Son*)). CRISPRtOE transposon insertion position and fluorescence measurements were determined for twelve isolates. Fluorescence measurements were normalized to cell density (OD_600_). Values are shown for isolates with on-target, RL orientation CRISPRtOE insertions and fold changes for on-target and RL orientation CRISPRtOE isolates compared to a strain with promoterless *mScarlet-I* (see Table S6 for details). Error is expressed for the median value of 8-12 isolates in n=3 (*Eco* and *Zmo*) or n=2 (*Ecl*, *Kpn*, *Aba*, *Pae*, *Ppu*, and *Son*) assays.

CRISPRtOE achieved a gradient of *mScarlet-I* OE across our series of tested promoters (Fig. 3B). CRISPRtOE containing the strong H promoter (P_H_) showed a ∼177-fold increase in mScarlet-I expression, while weaker promoters (P_V_ and P_A_) showed intermediate increases (∼2-30-fold); OE was validated at the RNA level using qPCR (Fig. S7A). CRISPRtOE lacking a promoter (no P) failed to stimulate expression above background, although we note that insertions in the reverse orientation (L-R) were excluded from this analysis due to cryptic promoter activity (Fig. S7B and Table S6). We also observed small variations in OE depending on the precise location of insertion for unknown reasons (Table S6).

We next sought to demonstrate CRISPRtOE activity in Proteobacteria with medical and industrial relevance. The non-model Alphaproteobacterium, *Zymomonas mobilis*, is a promising biofuel producer and an intriguing example of a free-living bacterium with a naturally reduced genome (37). To assay CRISPRtOE function in *Z. mobilis*, we again generated a promoterless *mScarlet-I* test strain and measured expression following insertion of CRISPRtOE transposons (Fig. S7C and Table S6). We found that CRISPRtOE could produce a gradient of overexpression in *Z. mobilis* up to ∼50-fold. Differences in the fold effects of CRISPRtOE across species likely reflect variation in promoter activity among our constitutive promoter set.

We then extended this approach to a panel of eight total species including the Gram-negative ESKAPE pathogens (38) *Klebsiella pneumoniae*, *Acinetobacter baumannii*, *Pseudomonas aeruginosa*, and *Enterobacter cloacae*, as well as the industrially and environmentally relevant strains *Pseudomonas putida* (39) and *Shewanella oneidensis* (40), focusing our strongest promoter (P_H_) to reduce experimental complexity (Fig. 3C and Table S6). We found that CRISPRtOE could increase mScarlet-I expression in all tested species with an OE range of 232-fold (*E. cloacae*) to 23-fold (*P. aeruginosa*), possibly due to variations in promoter activity, translation initiation or mScarlet-I folding across species. The CRISPRtOE insertion position in most species was centered at 33 bp upstream of *mScarlet-I*, although *A. baumannii* showed a wider range of positions for unknown reasons (Fig. S7D and Table S6). We note that the modular design of CRISPRtOE enables facile swapping of promoter sequences with organism-specific or inducible promoters to customize expression levels depending on the application and that we have constructed CRISPRtOE variants with a variety of antibiotic markers to facilitate its use in resistant environmental or clinical isolates. Recovery of CRISPRtOE transconjugants ranged from ∼10^-2^ to 10^-6^ across species (Table S7), with most species showing sufficient insertion efficiency to facilitate construction of genome-scale libraries. We conclude that CRISPRtOE is an effective overexpression strategy for Proteobacteria.

### Pooled CRISPRtOE Recapitulates Known Antibiotic Targets

There is an urgent need to develop new therapeutics to stem the rising tide of antibiotic resistance (41). One major bottleneck in the process of translating new antimicrobials to the clinic is determining the mode of action, including identifying the direct target. CRISPRtOE could provide a straightforward avenue to antibiotic target discovery because strains with overexpressed target proteins would outgrow competitor strains in a pooled context. For instance, it has been shown that OE of *folA*, encoding the trimethoprim (TMP) target, Dihydrofolate reductase (DHFR), can substantially increase resistance to TMP (42). Likewise, *murA*, encoding the fosfomycin (FOS) target, UDP-NAG enolpyruvyl transferase (MurA), can provide FOS resistance when overexpressed (43).

To validate CRISPRtOE as a tool for phenotyping antibiotic targets, we overexpressed the *folA* and *murA* genes in *E. coli* K-12 using the strong promoter construct (P_H_). As a hedge against potential variation in integration or OE efficacy, we tiled the region upstream of both genes with 12 unique spacers per gene and created a negative control library that inserted into the *lacZ* coding region (Fig. S8A). We then performed a pooled growth assay competing either the *folA* or *murA* library against the *lacZ* library in sub-lethal concentrations of TMP or FOS, respectively. Strikingly, we found that all CRISPRtOE gRNAs targeting *folA* and *murA* were effective in increasing antibiotic resistance (Fig. S8B and S8C). Relative to *lacZ* controls, the *folA* library grown in TMP showed a 22-fold median increase in abundance while the *murA* library grown in FOS showed a 52-fold median increase. To further characterize these resistance phenotypes, we performed Minimal Inhibitory Concentration (MIC) strip or broth microdilution assays on individual CRISPRtOE isolates (Fig. S8D, S8E, S8F, and S8G). We observed substantial shifts in MIC (∼15-30 fold) for CRISPRtOE insertions at varying distances upstream of *folA* (Fig. S8E and S8F), suggesting that CRISPRtOE phenotypes are robust to insertion distances upstream of target genes. We obtained similar results from measuring the FOS MIC for *murA* CRISPRtOE isolates (Fig. S8G).

Strong phenotypes from *folA* and *murA* OE provided an opportunity to test if CRISPRtOE targeting could be multiplexed. Thus, we targeted both *folA* and *murA* for OE in the same cell using two distinct strategies: 1) kanamycin (KAN)- or gentamicin (GEN)-marked transposons containing either a single *folA* or *murA* spacer were used in a quad-parental mating and transconjugants were selected on both KAN and GEN, or 2) a kanamycin-marked Tn*6677* containing a CRISPR array with both *folA* and *murA* spacers was used in a tri-parental mating and transconjugants were selected on KAN (Fig. S9A). We found that 97% of KAN^R^/GEN^R^ transconjugants from the quad-parental mating contained insertions upstream of both *folA* and *murA*, and we confirmed TMP^R^/FOS^R^ phenotypes for four tested isolates (Fig. S9B, S9C, and S9D). In contrast, only 12.5% of transconjugants from the tri-parental mating contained both *folA* and *murA* upstream insertions in the correct orientation; however, all four colonies tested were TMP^R^/FOS^R^ (Fig. S9B, S9C, and S9D). We conclude that CRISPRtOE is effective at targeting endogenous genes, such as antibiotic targets, and can be effectively multiplexed.

### Genome-scale CRISPRtOE Validates Antibiotic Targets and Reveals Resistance Pathways

Buoyed by our success with *folA* and *murA*, we expanded the scope of our CRISPRtOE screening approach to the genome-scale (Fig. 4A). We targeted all protein-coding genes in the *E. coli* K-12 genome with ∼10 gRNAs designed to integrate transposons within a small window upstream of genes without disrupting downstream coding sequences (∼45,000 guides total). Based solely on this design, we predicted that 64% of transposon insertions would occur in intergenic regions and 36% would occur in upstream genes (Fig. S10A and S10B). Thus, we generated two pooled libraries—one with promoter H and one without a defined promoter—to distinguish between phenotypes caused by OE rather than upstream gene disruption.

**Figure 4.**
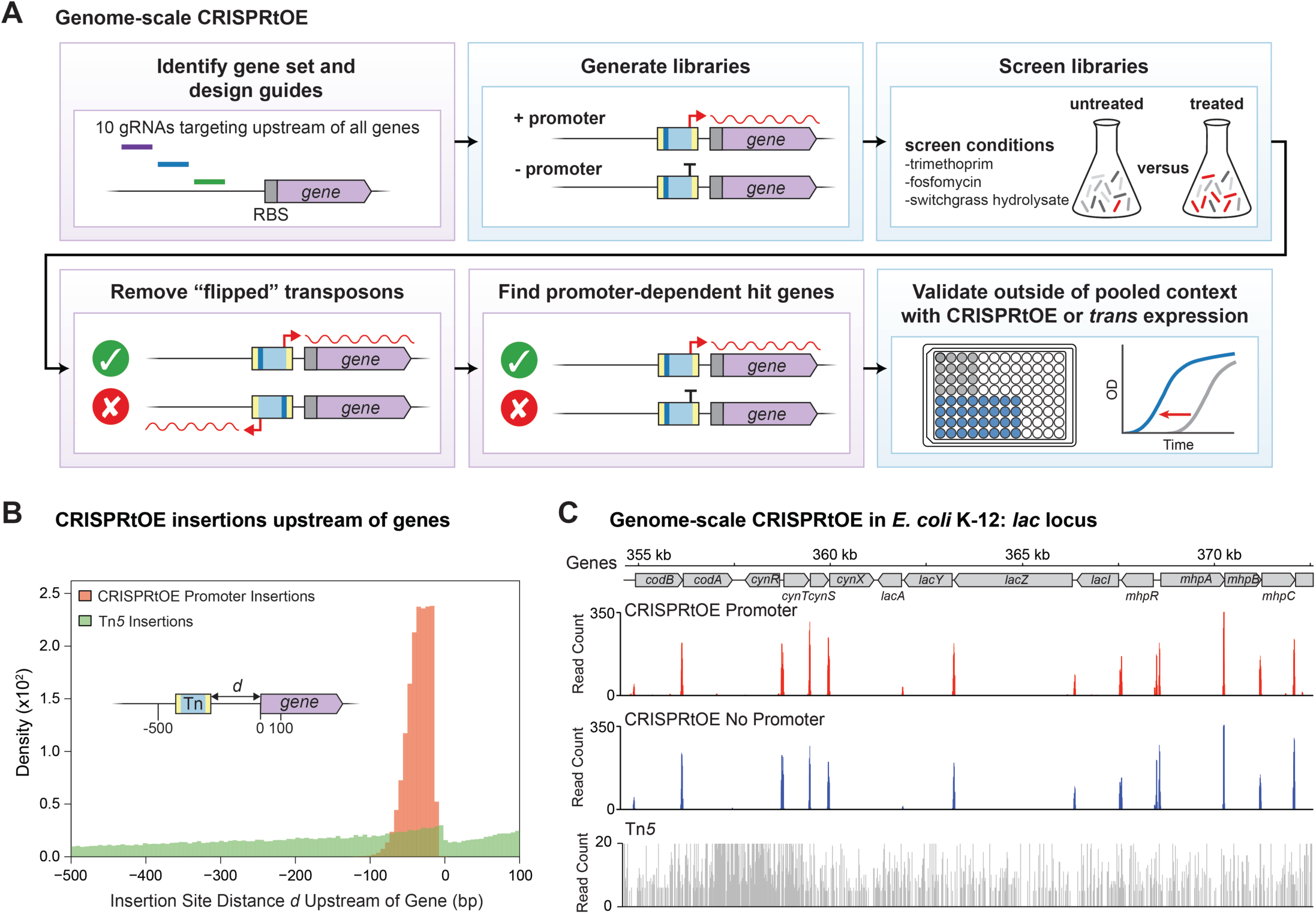
Genome-scale CRISPRtOE. (A) Schematic of genome-scale CRISPRtOE experiment including design and construction of libraries, experimental screen, bioinformatic data processing, and validation. Bioinformatic steps are in purple and experimental steps are in blue. Ten gRNAs were designed to guide insertion of the Tn6677 transposon (with or without a promoter) upstream of all genes. Pooled CRISPRtOE libraries grown in the presence of sub-lethal concentrations of ihibitors (e.g. *E. coli*: trimethoprim and fosfomycin antibiotics; *Z. mobilis*: switchgrass hydrolysate) and were screened for fitness by Tn-seq before and after treatment. Data were analyzed by counting reads that were in the R-L orientation upstream of genes to discover fitness phenotypes dependent on gene overexpression. Individual strains overexpressing genes with CRISPRtOE or from a plasmid were constructed to validate phenotypes outside of the pooled context. (B) Positions of CRISPRtOE insertions upstream of all genes in the genome. Insertion site distance d was calculated based on the distance in bp from the 3′ end of the Tn to the 5′ end of the gene. CRISPRtOE Tn insertions shown in red. For comparison, random/untargeted insertions from a Tn*5* library are shown in green. (C) CRISPRtOE insertions into the *E. coli lac* locus showing precise placement upstream of genes compared to a library constructed with Tn*5* random transposition. The scale is capped at 20 reads for visual clarity.

We found that CRISPRtOE effectively targeted regions upstream of genes across the genome. We first used Tn-seq to map transposon locations, then performed a computational analysis to associate insertions in the proper orientation (R-L) with downstream genes. Based on this analysis, an average of ∼97% of insertions occurred at locations consistent with the library design, demonstrating high, on target efficiency (Fig. S10C, Table S8). Unfortunately, transposon insertion locations could not be defined using gRNA spacer sequences alone, as a fraction of transconjugant cells likely obtained and expressed multiple gRNAs during the mating procedure. This resulted in transposons that inserted at a site designated by the library design that nonetheless did not match the gRNA spacer found within the inserted transposon (“hijacked” transposons). Although our libraries showed that, on average, ∼29% of transposon insertions were hijacked (Fig. S10C, Table S8), we estimate that only ∼3.5% of transconjugants in our library contained more than one insertion based on mating experiments that used differentially marked transposons (Fig. S10D). In our Tn-seq analysis, we found that CRISPRtOE insertions occurred in a tight distribution largely within 100 bp of the 5′ ends of genes (Fig. 4B). This is in stark contrast to the nearly flat distribution of Tn*5* insertions, which are known to occur pseudo-randomly (44). For instance, the visual distinction between CRISPRtOE targeted and Tn*5* pseudo-random insertion was readily apparent at the non-essential *lac* locus (Fig. 4C).

To test the ability of CRISPRtOE to identify antibiotic-relevant gene phenotypes at the genome-scale, we grew our libraries in the presence or absence of a sublethal concentration of TMP and measured relative strain abundance. Biological replicates showed excellent agreement at the gene level (R^2^ ∼ 0.92, Fig. S11A), underscoring the high reproducibility of genome-scale CRISPRtOE. The *folA* gene was a clear positive outlier with >1000-fold enrichment in TMP versus untreated (Fig. 5A and 5B), demonstrating the power of our CRISPRtOE screen in recovering antibiotic targets. Substantial *folA* enrichment was only seen in the P_H_-containing library and not in the promoterless library, indicating that OE, rather than transposon insertion per se, was responsible for the phenotype (Fig. 5B). Promoter-dependent phenotypes were largely distinct from insertion phenotypes across the entire genome, as relative strain abundances of the P_H_ and no promoter libraries after TMP treatment were not correlated (R^2^ = 0.007, compare Fig. S11B and S11C). Computational subsampling of our libraries suggested that CRISPRtOE library complexity could be reduced, if desired. Gene-level phenotypes remained robust after subsampling the library from ∼10 to 4 random guides per gene (R^2^ = 0.905, Fig. S12A). Further, we obtained many of the same top hits, including *folA*, if we only considered guides that targeted the first gene in operons (Fig. S13A). Thus, we anticipate that future CRISPRtOE libraries could achieve genome-scale overexpression with far fewer guides per gene.

**Figure 5.**
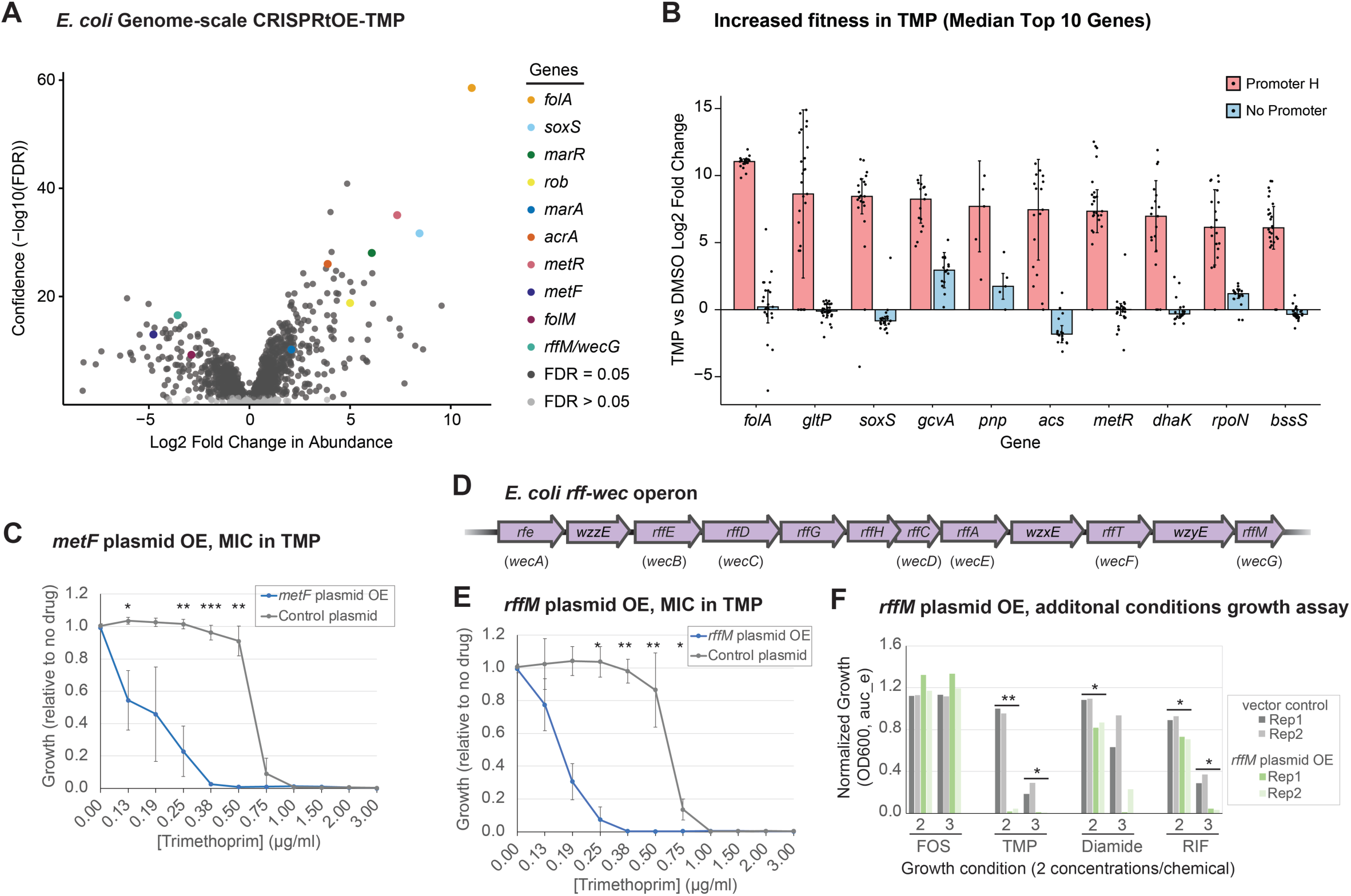
*E. coli* genome scale CRISPRtOE screen with trimethoprim. (A) Volcano plot of gRNA spacer counts in trimethoprim (TMP) versus dimethylsulfoxide (DMSO) control. Each dot represents a strain with a spacer targeting upstream of a gene. Those that are not statistically significant (false discovery rate (FDR) > 0.05) are shown in light grey and significant hits (FDR ≤ 0.05) are shown in dark gray. Several outliers are colored: e.g. *folA* (red), *metF* (blue), *rffM/wecG* (green). (B) Top 10 gene hits with increased fitness in TMP treatment. Genes must have at least a 4-fold change in the CRISPRtOE-Promoter H data and show a 4-fold difference in a comparison between Promoter H (red bars) and no promoter (blue bars) to be considered. Dots represent unique insertions. Error bars represent standard deviation (SD). (C) Liquid MIC assay showing resistance of strains overexpressing *metF* from a plasmid to TMP. Error bars represent SD of triplicate assays. Significance was determined by a t test (≤0.05(*), ≤0.01(**), ≤0.001(***)). (D) *E. coli* Enterobacterial Common Antigen (ECA) biosynthesis (*rff*/*wec*) operon. (D) Liquid MIC assay showing resistance of strains overexpressing *rffM* from a plasmid to TMP. Error bars represent SD of triplicate assays. Significance was determined by a t test (≤0.05(*), ≤0.01(**), ≤0.001(***)). (F) Growth assay of strains overexpressing *rffM* from a plasmid in additional chemical conditions (Biolog phenotypic array plate PM16 comprised of 24 compounds). Two concentrations of CRISPRtOE screen chemicals FOS (no response expected) and TMP (response expected), and of two additional chemical that had a significant response (diamide and rifamycin) are shown. Duplicate assays are shown (Rep1 and Rep2). Significance was determined by a t test (≤0.05(*), ≤0.01(**). See figure S14 for additional details.

Significant outliers outside of the direct target revealed potential TMP resistance or susceptibility mechanisms (Fig. 5A and 5B). For instance, one of the most resistant outlier genes was *soxS*, encoding an AraC-type transcription activator known to regulate genes involved in antibiotic resistance (45) (Fig. 5B). Indeed, AraC and MarR family transcription factor genes and their regulons were among the most resistant outliers in our screen (e.g., *marR*, *rob*, *marA* and the SoxS/Rob/MarA regulated AcrAB efflux pump (46), Table S9), highlighting the ability of CRISPRtOE to identify resistance gene regulators by amplifying natural activation circuits.

Other resistance and susceptibility mechanisms centered on folate metabolism. TMP treatment reduces cellular methionine levels due to the involvement of tetrahydrofolate (the product of DHFR) in methionine synthesis (47). CRISPRtOE of the *metR* gene, which encodes a positive regulator of methionine biosynthesis genes (48), increased TMP resistance (Fig. 5B), possibly in increasing flux through the pathway. In contrast, *metF* OE was highly deleterious in the presence of TMP (Fig. 5A), likely due to depletion of 5,10-methylenetetrahydrofolate pools normally used by the ThyA-DHFR pathway to generate tetrahydrofolate (49). We validated this toxicity by showing that *metF* OE from a multi-copy plasmid reduces the MIC for TMP by ∼4-fold (Fig. 5C). Neither *metR* nor *metF* were recovered in previous TMP loss-of-function screens (50, 51), demonstrating the utility of CRISPRtOE to identify new players in antibiotic function. Interestingly, CRISPRtOE of *folM*, which is thought to encode a protein with DHFR activity (52), was highly toxic to TMP-treated cells (Fig. 5A, Table S9). Although unintuitive, this result is consistent with other genome-scale phenotyping studies that found that disruption of *folM* increases resistance to TMP (50, 51).

Finally, genes involved in cellular polysaccharide synthesis, including Enteric Common Antigen (ECA) biosynthesis genes (e.g., genes in the *rff*/*wec* operon), were functionally enriched among sensitive outliers (FDR = 0.0054, ks test; Table S10A). Previous work has shown that disruption of the ECA pathway also increases TMP sensitivity, possibly by sequestering thymidine pools and leading to “hyper-acting” version of thymine-less death (53). Our work suggests that dysregulation by OE, in addition to disruption, can cause ECA-dependent toxicity in TMP-treated cells. Although ECA genes exist in complex operons that may be disrupted by insertion (Fig. 5D), phenotypes for some ECA genes required the presence of P_H_ in the transposon (Table S9). We chose to focus our follow up analysis on *rffM* (a.k.a. *wecG*), which is the last gene in the ECA operon and showed a phenotype that was strongly dependent on P_H_ (Table S9). To validate TMP sensitivity upon *rffM* OE and to rule out *cis* effects from disruption of the ECA operon, we expressed *rffM* from a multi-copy plasmid. Plasmid-based OE of *rffM* reduced the TMP MIC by ∼4-fold (Fig. 5E), confirming that OE is the source of toxicity. As ECA is found on the cell surface, we considered that ECA pathway disruption by *rffM* OE might increase susceptibility to a broader set of small molecule inhibitors other than TMP. To test this, we screened our plasmid-based *rffM* OE strain against 24 chemicals using a Biolog Phenotype Array. We found that *rffM* OE increased sensitivity to additional small molecule compounds, such as diamide (a.k.a., tetramethylazodicarboxamide) and rifamycin SV (Fig. 5F and S14A and S14B). Thus, OE of ECA synthesis genes increases susceptibility to TMP and other small molecule inhibitors, although the precise mechanism and scope of affected inhibitors remains unknown.

We next expanded our genome-scale proof-of-principle studies by screening our CRISPRtOE library against FOS, finding expected, underexplored, and potentially novel FOS resistance determinants (Fig. 6 and S15 and Table S11). As expected, *murA* was a prominent resistant outlier along with the gene upstream of *murA*, *ibaG* (Fig. 6A and S15A). As OE of *ibaG* and *murA* showed similar fold increases in FOS treated cells, we speculated that the *ibaG* phenotype is due to increased transcription of *murA* rather than a contribution of the *ibaG* gene product to FOS resistance, per se.

**Figure 6.**
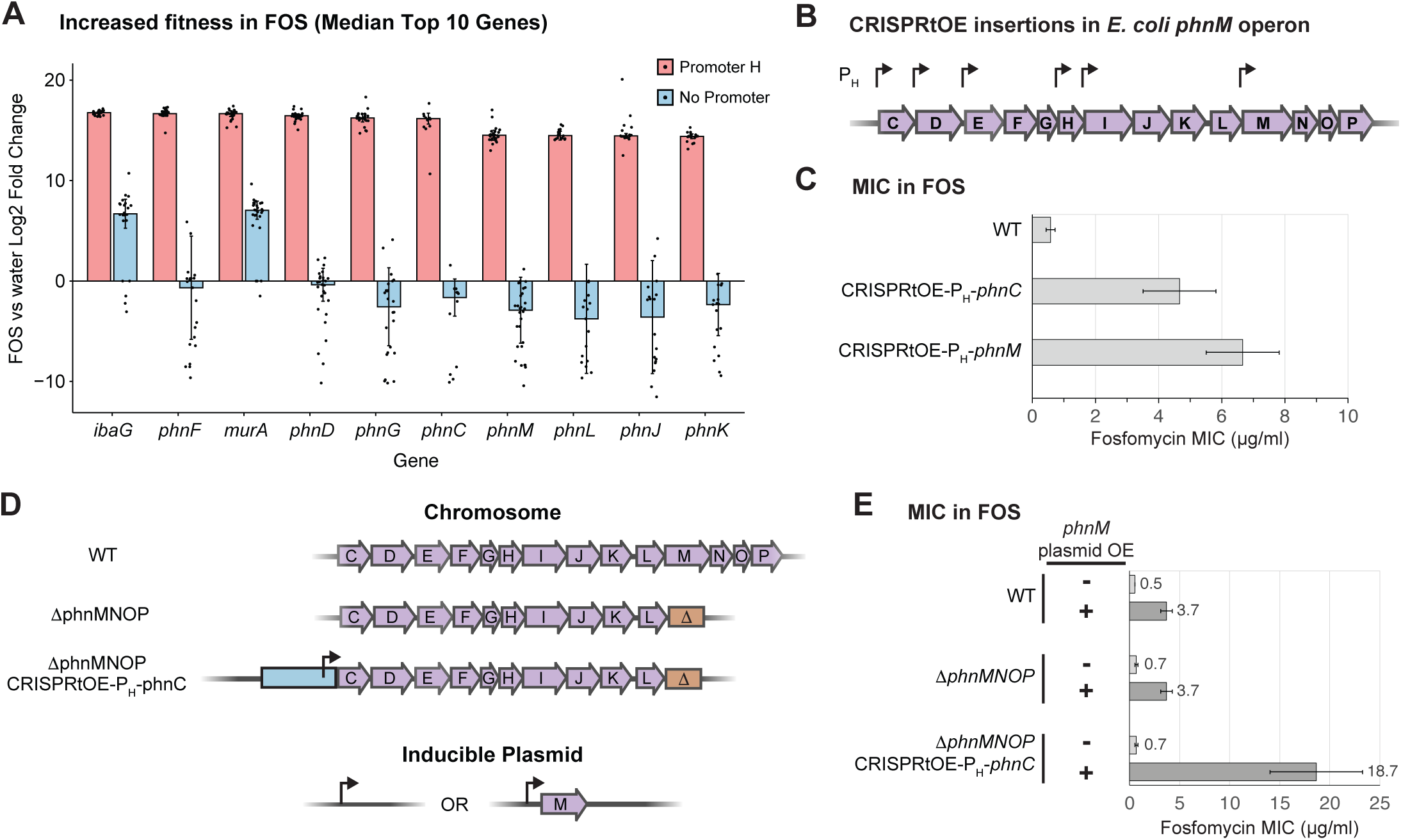
*E. coli* genome scale CRISPRtOE screen with fosfomycin. (A) Top gene hits with increased fitness in fosfomycin (FOS) treatment. Genes must have at least a 4-fold change in the CRISPRtOE-Promoter H data and show a 4-fold difference in a comparison between Promoter H (red bars) and no promoter (blue bars) to be considered. Dots represent unique insertions. sError bars represent standard deviation (SD). (B) CRISPRtOE-Promoter H insertions with increased fitness in FOS in the *E. coli* phosphonate utilization (*phn*) operon represented by arrows. (C) Quantification of MIC test strip (Liofilchem) assay showing resistance to FOS of individual CRISPRtOE-Promoter H strains overexpressing *phnC* or *phnH*. Error bars represent SD of triplicate assays (also see figure S15B). (D) Schematic of *phnMNOP* chromosomal deletion and *phnM* complementation experiment. (E) Quantification of FOS MIC test strip (Liofilchem) assays in WT, *ΔphnMNOP* deletion, or CRISPRtOE-Promoter H-*phnC* plus *ΔphnMNOP* deletion strains that are overexpressing *phnM* from a plasmid (or empty vector control). Error bars represent SD of triplicate assays (also see figure S15D).

Among the other top resistance genes, we identified several genes within the *phn* operon (*phnCDEFGHIJKLMNOP*) (Fig. 6A and S15A and Table S11) which is involved in degradation of small molecules containing carbon-phosphate (C-P) bonds known as phosphonates (54). Intriguingly, FOS contains a C-P bond that is central to the molecule and likely critical for its function as an antibiotic. The *phn* operon was previously implicated in FOS resistance in a screen that used randomly inserting transposons that contained promoters (55), but the genetic basis of the OE phenotype was not further characterized. We found that CRISPRtOE insertions upstream, but not downstream of *phnM* allowed for FOS resistance, suggesting that OE of *phnM*, *phnN*, *phnO*, or *phnP* was responsible for the phenotype (Fig. 6B). We first validated *phnC* and *phnM* CRISPRtOE phenotypes outside of the pooled screen context, finding increases in the FOS MIC of ∼8- and ∼11-fold, respectively (Fig. 6C and S15B). Deletion of *phnMNOP* had no impact on the FOS MIC of otherwise wild-type (WT) cells, demonstrating that the role of *phn* genes in FOS resistance is only apparent through OE (Fig. 6D, 6E, and S15C). However, deletion of *phnMNOP* eliminated the increase in FOS resistance caused by *phnC* CRISPRtOE (Fig. 6E and S15D). Reasoning from the insertion data that OE of *phnM* was most likely to be the key player in our FOS resistance phenotype, we overexpressed *phnM* from a multi-copy plasmid and measured FOS MICs. Consistent with our hypothesis, we found that *phnM* OE was sufficient to increase the FOS MIC in otherwise wild-type cells and for a strain lacking *phnMNOP* on the chromosome (Fig. 6E and S15D). Interestingly, plasmid-based *phnM* OE in a *phnC* CRISPRtOE Δ*phnMNOP* background further increased the FOS MIC to ∼37-fold over WT (Figures 5E and S15D), suggesting that *phnM* functions alongside other members of the *phn* operon, but also acts as a critical bottleneck to FOS resistance in WT cells.

Finally, we investigated the role of *waaY*, which encodes the lipopolysaccharide (LPS) core heptose II kinase (56), in FOS resistance. In the absence of *waaY*, LPS is not phosphorylated on heptose II (56) and cells become sensitized to a number of small molecule inhibitors and antibiotics, but not FOS (45). However, our CRISPRtOE screen suggested that *waaY* OE could provide resistance to FOS (Fig. S15A and S15E). We confirmed that *waaY* increases the FOS MIC when overexpressed in *cis* (via CRISPRtOE) or *trans* (via multi-copy plasmid). Thus, our experiments suggest that cellular levels of *waaY* and by extension, LPS phosphorylation, are insufficient to provide FOS resistance but OE can overcome this deficit. In sum, genome-scale CRISPRtOE recovered known antibiotic targets and resistance mechanisms as well as intriguing new players in resistance.

### Genome-scale CRISPRtOE in a non-model bacterium identifies potential engineering targets

We next sought to extend our genome-scale CRISPRtOE approach beyond model bacteria and well-characterized inhibitors. *Zymomonas mobilis* is an Alphaproteobacterium that shows promise in conversion of deconstructed plant material (hydrolysate) into biofuels. Plant hydrolysates contain sugars used by *Z. mobilis* in the production of biofuels, but also a complex mixture of inhibitory compounds that originate from the deconstruction process or from the plant material itself. Only some of these inhibitors are known. In principle, OE screens could identify engineering targets that overcome the growth defects caused by hydrolysate inhibitors and increase biofuel yields. To identify potential engineering targets, we constructed CRISPRtOE libraries containing no promoter or P_H_ that targeted all protein coding genes in the *Z. mobilis* ZM4 genome (∼20,000 guides total) using the same design principles described above for *E. coli*. We used Tn-seq to map transposon locations and found high on-target efficiency (Fig. S10C). We then screened our libraries against a concentration of ammonia fiber expansion (AFEX)-treated switchgrass hydrolysate that resulted in ∼50% growth inhibition when added to defined medium. We found multiple genes that improved the relative and absolute growth of *Z. mobilis* in switchgrass hydrolysate (Fig. 7A and 7B and Table S12). We focused our follow up studies on two intriguing resistant outliers, *fumA*, encoding Fumarase A, and *rpoE*, encoding an extracytoplasmic function (ECF) σ factor (Fig. 7, S16, and S17). Fumarase typically participates in the tricarboxylic acid (TCA) cycle, but *Z. mobilis* has an incomplete TCA pathway lacking key enzymes such as malate dehydrogenase (57). CRISPRtOE and plasmid-based OE of *fumA* outside the pooled context validated the growth advantage of *fumA* over WT, although the absolute magnitude of this advantage was modest (Fig. S16A, S16B, S17A, and S17B). However, CRISPRtOE and plasmid-based OE of *rpoE* in hydrolysate showed a much larger absolute growth effect versus WT (Fig. 7C, 7D, S16C, S16D, S17C, and S17D). As many ECF σ factors are involved in mitigating cell stress, we speculate that *rpoE* OE “pre-induces” a cellular stress response which reduces the long lag phase associated with growth in hydrolysate.

**Figure 7.**
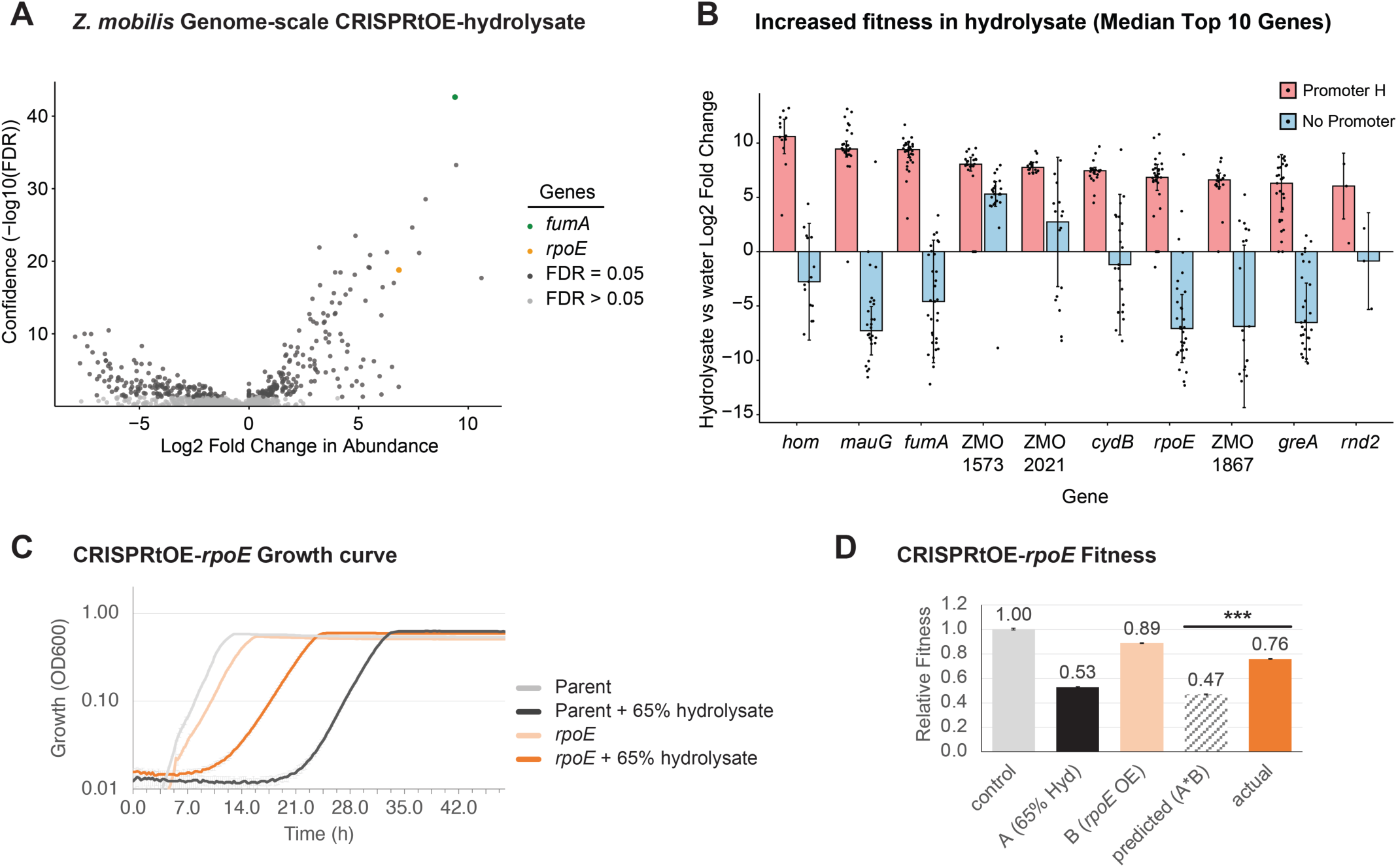
*Z. mobilis* genome scale CRISPRtOE screen with plant hydrolysate. (A) Volcano plot of gRNA spacer counts in 65% hydrolysate versus *Zymomonas* rich defined medium (ZRDM) control. Each dot represents a strain with a spacer targeting upstream of a gene. Those that are not statistically significant (false discovery rate (FDR) > 0.05) are shown in light grey and significant hits (FDR ≤ 0.05) are shown in dark gray. Several outliers are colored: *fumA* (green), *rpoE* (orange). (B) Top gene hits with increased fitness in 65% hydrolysate. Genes must have at least a 4-fold change in the CRISPRtOE-Promoter H data and show a 4-fold difference in a comparison between Promoter H (red bars) and no promoter (blue bars) to be considered. Dots represent unique insertions. Error bars represent standard deviation (SD). (C) Growth assay of individually constructed CRISPRtOE-Promoter H *rpoE* strains versus the parent strain in 65% hydrolysate or ZRDM control. Error bars represent standard deviation (SD) of three independant cultures. (D) Relative fitness of strains shown in panel (C). Growth was quantified by calculating the area under the curve (auc_e) using GrowthCurver. Fitness was normalized to control condition (parent strain in ZRDM). Significance was determined by a t test (≤0.005(***). See also Figure S16C and S16D.

However, we caution that very little is known about the RpoE regulon in *Z. mobilis*, and the name “*rpoE*” may be misleading as the *Z. mobilis* protein shows low percent identity with well-characterized RpoE proteins from other bacteria (e.g., *E. coli* K-12 RpoE (58) (b2573) ∼23%, *Rhodobacter sphaeroides* RpoE (59) (RSP_2681) ∼40%). In sum, we extended genome-scale CRISPRtOE to the non-model bacterium *Z. mobilis* and successfully identified genes that improved growth in complex plant hydrolysate.

## Discussion

Functional genomics approaches that are robustly scalable and readily applicable to diverse bacteria are required to bridge the yawning gap between genome sequencing and gene function assessment. Our work provides a targeted, systematic, and practical approach for genome-scale gene overexpression in Proteobacteria and possibly beyond. By modifying VcCAST for use in functional genomics (Fig. 1 and 2) and defining non-transcribed regions of DNA as high-efficiency insertion sites (Fig. S3), we demonstrated targeted and tunable OE in Proteobacteria with medical, industrial, and basic research relevance (Fig. 3). Our genome-scale OE screen for targets and modulators of TMP and FOS efficacy in the model organism *E. coli* highlight the exquisite specificity of CRISPRtOE in recovering genes and pathways that underpin antibiotic function (Fig. 4, 5 and 6). The extension of our genome-scale OE screening approach to the non-model bacterium *Z. mobilis* in a complex inhibitor mixture and the ease of constructing individual CRISPRtOE inhibitor-resistant strains demonstrates the potential utility for industrial strain optimization (Fig. 7). Given the ease of implementing CRISPRtOE in the organisms explored here, we anticipate that our approach will be readily expandable to many other microbes and screening conditions. While our manuscript was under review, another manuscript was published that showed that VcCAST containing promoters could be used to upregulate genes in *E. coli*(32). Our work dramatically expands this approach across multiple Proteobacteria and to the genome scale. We note that the replicative CAST vector used in that study is suboptimal for use at the genome-scale due to a lack of Tn selection, the need to cure the plasmid post-editing, and an inability to replicate in important genera such as *Acinetobacter*.

CRISPRtOE is a valuable genetic tool for both basic and applied biology. The targeted aspect of CRISPRtOE can be used to generate focused libraries that limit size (number of gRNAs) and scope (number of targeted genes). Moreover, screen hits can be readily pursued for downstream mechanistic analysis by individually cloning gRNAs used in the screen to easily recreate strains outside of the pooled context, a clear benefit over random transposition approaches such as Tn*5*. Additionally, generating pooled sub-libraries from screen hits enables follow up validation at scale. Importantly, the ability to insert transposons with or without promoters at the same genomic locus will allow researchers to disentangle the effects of Tn insertion versus overexpression, simplifying hit interpretation. Further, combining CRISPRtOE with existing gene perturbation approaches such as CRISPRi could enable screening for overexpression suppressors of essential functions at the genome-scale.

Our proof of principle work demonstrating that genome-scale CRISPRtOE can identify resistance genes in *E. coli* K-12 holds promise for extending this approach to clinically relevant pathogens with the goal of improving diagnosis and treatment. Importantly, CRISPRtOE action is similar to clinically relevant antibiotic resistance mechanisms, such as IS element transposition and overexpression of downstream resistance genes (60). CRISPRtOE may also be valuable for optimization of strains for industrial use. Industrial strain optimization often occurs through loss of function (61) or directed evolution experiments (62), followed up by plasmid-based overexpression of mutated pathways. CRISPRtOE streamlines this process in non-model bacteria, particularly by enabling recreation of optimized strain features in different genetic backgrounds by targeted transposition. Finally, CRISPRtOE-optimized strains may be immediately useful in production, as CRISPRtOE leaves no heterologously expressed *cas* genes in the recipient strain.

Several open questions remain regarding the mechanisms by which genes found in our CRISPRtOE screens provide resistance to TMP, FOS, and hydrolysate. Do potential alterations in the outer membrane caused by *rffM* OE alter cell permeability to small molecule inhibitors such as TMP? RffM catalyzes the first committed step in the pathway to generate ECA modified LPS (ECA-LPS); thus, *rffM* OE would be predicted to increase the ratio of ECA-LPS to N-acetyl-D-glucosamine modified LPS (GlcNAc-LPS) in *E. coli* K-12 (or O-antigen modified LPS in other *E. coli* strains). Consistent with this the Bernhardt lab recently showed that *rffM* OE dramatically reduced GlcNAc-LPS-specific labeling (63), demonstrating perturbation of the outer membrane. Does PhnM act in concert with other members of the Phn enzymatic machinery to modify and degrade FOS? Phn operon members are thought to degrade phosphonates by the following mechanism: *i*) PhnI along with PhnG, PhnH, and PhnL use ATP and phosphonate compounds as substrates to produce 5-triphosphoribosyl-α-1-phosphonate, which is then *ii*) degraded by PhnM into pyrophosphate and 5-phosphoribosyl-α-1-phosphonate, and finally *iii*) the C-P bond is broken by PhnJ as part of the large Phn(GHIJKL)_2_ complex (64, 65). PhnM lacks C-P lyase activity and is unlikely to directly degrade FOS; however, it is possible that PhnM-mediated conversion of the triphosphorylated phosphonate substrate into a monophosphorylated form is limiting flux through the pathway. Alternatively, if PhnM is sufficient for FOS resistance, by what mechanism does it act? Does the extent of WaaY-dependent, heptose II phosphorylation of LPS mirror the extent of resistance provided by *waaY* OE? Although WaaY appears to be required for phosphorylation (56), the fraction of modified LPS molecules in WT versus *waaY* OE is unknown. Finally, what target genes are induced upon *rpoE* OE to drive resistance to hydrolysate toxins? Mutations in the transcription machinery often occur during laboratory evolution experiments to optimize growth in complex or pleiotropic conditions (66), suggesting that the expression of multiple genes needs to be altered to satisfy the genetic selection. Future work will map the RpoE regulon in *Z. mobilis*, potentially identifying additional stress resilience factors and permitting a comparison to better characterized ECF σs.

CRISPRtOE is not without caveats and there may be scenarios in which it is outperformed by other overexpression or gene manipulation strategies. CRISPRtOE and other engineering approaches that insert promoters will alter expression of downstream genes in operons; these effects could complicate but not necessarily preclude identification of gene-level CRISPRtOE phenotypes. Our view is that expression of downstream genes is often advantageous due to fact that protein complexes are often found in operons (e.g., efflux pump subunits). We expect that “No promoter” CRISPRtOE libraries will largely allow for disambiguation of insertion versus overexpression phenotypes, but there may be rare instances in which the combination of insertion and overexpression causes an unexpected genetic interaction because operons contain functionally related genes. In this scenario, Dub-seq or promoter recombineering strategies would be advantageous, as Dub-seq expresses genes *in trans* and recombineering can be used to introduce more subtle *cis* mutations that could avoid perturbing upstream genes. Dub-seq can also introduce heterologous DNA from other species for phenotyping (14), something CRISPRtOE is not designed to accomplish. Although we have only demonstrated genome-scale CRISPRtOE in selected Proteobacteria, we envision that improvements to the technology by ourselves and others will enable broad use of the technology in diverse bacteria.

## Materials and Methods

### Strains and growth conditions

Strains are listed in Table S1. Growth and antibiotic selection conditions are summarized in Table S2. *Escherichia coli*, *Acinetobacter baumannii*, *Enterobacter cloacae*, *Klebsiella pneumoniae,* and *Pseudomonas aeruginosa* were grown in LB broth, Lennox (BD240230) at 37°C (or 30°C for CRISPR-guided transposition) in a flask with shaking at 250 rpm, in a culture tube on a roller drum, in a 96 well deepwell plate with shaking at 900 rpm, or in a Tecan Sunrise plate reader shaking with OD600 measurements every 15 min. *Pseudomonas putida* and *Shewanella oneidensis* were grown in LB in a culture tube on a roller drum or in a 96 well deepwell plate with shaking at 900 rpm at 30°C. *Zymomonas mobilis* was grown in RMG medium (10g yeast extract and 2g KH_2_PO_4_ monobasic/liter with 2% glucose added after autoclaving) at 30°C statically in a culture tube or deepwell plate. *E. coli* was grown in Mueller Hinton Broth (MHB, BD 275730) for antibiotic minimal inhibitory concentration (MIC) assays. Media was solidified with 1.5-2% agar for growth on plates. Antibiotics were added when necessary: *E. coli* (100 µg/ml carbenicillin (carb), 30 µg/ml kanamycin (kan), or 50 µg/ml spectinomycin (spt)), *A. baumannii* (60 µg/ml kan or 100 µg/ml apramycin (apr)), *E. cloacae* (30 µg/ml kan or 50 µg/ml spt), *K. pneumoniae* (60 µg/ml kan or 50 µg/ml spt), *P. aeruginosa* (30 µg/ml gentamicin (gen) or 1 mg/ml kan), *P. putida* (150 µg/ml apr or 60 µg/ml kan), *S. oneidensis* (60 µg/ml kan or 30 µg/ml gen), and *Z. mobilis* (200 µg/ml gen or 100 µg/ml spt). Diaminopimelic acid (DAP) was added at 300 µM to support growth of *dap-E. coli* strains. Strains were preserved in 15% glycerol at -80°C.

### General molecular biology techniques

Plasmids are listed in Table S3A and a summary of plasmids for utilizing the CRISPRt and CRISPtOE systems and constructing mScarlet-I reporter strains is in Table S3B. Oligonucleotides are listed in Table S4A (individual) and Tables S4B-E (pooled). *pir*-dependent plasmids were propagated in *E. coli* strain BW25141 (sJMP146) or its derivative sJMP3053. Plasmids were purified using the GeneJet Plasmid Miniprep kit (Thermo K0503), the QIAprep Spin Miniprep Kit (Qiagen 27106), or the Purelink HiPure Plasmid Midiprep kit (Invitrogen K210005). Plasmids were digested with restriction enzymes from New England Biolabs (NEB, Ipswich, MA). Ligations used T4 DNA ligase (NEB M0202) and fragment assembly used NEBuilder Hifi (NEB E2621). Genomic DNA was extracted with the DNeasy Blood & Tissue Kit (Qiagen 69504) or the GeneJet genomic DNA purification kit (Thermo K0721). DNA fragments were amplified by PCR using Q5 DNA polymerase (NEB M0491, for cloning) or One*Taq* DNA Polymerase (NEB M0480, for analysis). PCR products were spin-purified using the Monarch PCR & DNA Cleanup Kit (NEB T1030) or the DNA Clean & Concentrator-5 kit (Zymo Research, Irvine, CA, D4013). Reactions were purified with 1.8X Mag-Bind TotalPure NGS magnetic beads (Omega) on a magnetic rack (Alpaqua). DNA was quantified spectrophometrically using a NanoDrop Lite or fluorometrically using a Qubit with the HS or BR DNA kit (Thermo). Plasmids were transformed into electrocompetent *E. coli* cells using a 0.1 cm cuvette (Fisher FB101) and a BioRad Gene Pulser Xcell on the Bacterial 1 *E. coli* preset protocol (25 µF, 200 ohm, 1800 V) as described in detail previously(67). Oligonucleotides were synthesized by Integrated DNA Technologies (Coralville, IA) or Agilent (SurePrint Oligonucleotide library) (Santa Clara, CA). Sequencing was performed by Functional Biosciences (Madison, WI), Plasmidsaurus (Eugene, OR), Azenta (South Plainfield, NJ), or the University of Wisconsin-Madison Biotechnology Center Next Generation Sequencing Core (UWBC NGS core, Madison, WI).

### *E. coli* Δ*lac* strain construction

An *E. coli* MG1655 strain with a *lac* operon deletion (Δ*lac*, Lac^-^) was constructed by λ-Red-mediated recombination as previously described(68). Briefly, *E. coli* MG1655 (sJMP163) harboring the plasmid pSIM6 (pJMP170) encoding the λ-Red proteins was transformed with a linear DNA fragment encoding an FRT-*kan*-FRT cassette which was amplified by PCR from plasmid pJMP3099 using primers oJMP912 and oJMP913 which also have 40 nt homology to the chromosomal insertion site) resulting in strain sJMP3267. The allele was transferred from sJMP3267 to *E. coli* MG1655 (sJMP163) by P1–*vir*-mediated transduction, as previously described(69), resulting in strain sJMP3269. The *kan* gene was removed by transformation with plasmid pCP20 (pJMP3008), encoding a constitutively expressed FLP recombinase, as previously described(70), resulting in strain sJMP3272. Full deletion of the *lac* operon in sJMP3272 was confirmed by PCR with flanking primers (oJMP914 and oJMP915) followed by Sanger sequencing.

### Construction of chromosomal fluorescent reporters (insertion into the *att*_Tn*7*_ site)

Fluorescent reporter cassettes are located in a Tn*7* transposon located on an R6K ori (*pir* dependent) plasmid. The transposons were transferred to the *att*_Tn*7*_ site ∼50 bp downstream of *glmS* of a recipient bacterial chromosome by co-conjugation of the Tn*7* transposon donor strain and a second donor strain harboring a plasmid expressing Tn*7* transposase(71). Donor strains encode conjugation machinery, encode *pir*, and are DAP dependent. Briefly, a donor strain with the transposon plasmid, a second donor with the transposase expression plasmid, and the recipient strain were resuspended from plates to an OD_600_=9 and mixed in equal proportions (100 µl each). Cells were harvested by centrifugation at 5000 rcf for 2 min (except *A. baumannii* at 9000 rcf) and placed on a nitrocellulose filter (Millipore HAWP01300) on an LB plate (except *Z. mobilis* on RMG) and incubated at 30°C (*Z. mobilis* and *P. putida*) or 37°C (all other organisms) for 2-18h. Cells from the filter were resuspended in 300 µl media, serially diluted, and plated with antibiotic selection for the transposon (see growth and selection conditions above) and without DAP to select against the donor strains, and incubated at 30 or 37°C until colonies formed.

A transposon containing a GFP expression cassette was derived from a Mobile-CRISPRi plasmid(67) by assembling the AscI-EcoRI fragment of pJMP2824 and annealed oligonucleotides oJMP1420 and oJMP1421 to create pJMP6957. The GFP encoding gene of pJMP6957 was replaced by an mScarlet-I encoding gene derived from Addgene plasmid 85069 (pJMP7001) to create pJMP10180 which has an *mScarlet-I* cassette (BbsI sites removed and *mScarlet* converted to *mScarlet-I*) with the T7A1 constitutive promoter in the Tn*7* transposon. A promoterless *mScarlet-I* reporter with the CRISPRt *LZ1* protospacer was derived from pJMP6957, pJMP10180, and synthetic DNA oJMP1599 to create pJMP10206 (see Table S3A for details).

### Construction of CRISPRt/CRISPRtOE system

The CRISPRt/CRISPRtOE donor plasmids were derived from pSpinR (pSL1765)(21), Mobile-CRISPRi plasmids(71), and synthetic DNA. Cloning details are located in Table S3A for the individual plasmids. In summary, (1) a functional antibiotic resistance cassette was added to the transposon (pJMP747) and the BsaI cloning site in the gRNA cassette was altered (pJMP10009 and pJMP10011). A two plasmid R6K ori (*pir*-dependent) system for selectable CRISPR-guided transposition was created by combining (1) the TnsABC-TniQ-Cas876 expression cassette from pSpinR with the amp^R^, R6K ori, mobilizable backbone from Mobile-CRISPRi (pJMP2782) to create pCRISPRt-H (pJMP10233) and (2) the transposon and guide RNA cassette from pJMP10011 with the same backbone (from pJMP2782) to create pCt-T (pJMP10237 and derivatives) which was further minimized, optimized, and altered by adding transcription terminators and moving the guide RNA cassette inside the transposon to produce pCRISPRt-T-gent (pJMP10395) and pCRISPRt-T-kan (pJMP10397). CRISPRtOE plasmids with either no promoter or various strength promoters (pCtOE-noP, pCtOE-P_A_, pCtOE-P_H_, pCtOE-P_V_) were derived from these plasmids by inserting a transcription terminator and promoter cassette derived from synthetic DNA (oJMP2000 plus oJMP2001-2004) into the above plasmids to produce plasmids oJMP10471-10479. Apramycin resistant derivatives of some of these plasmids were created by digesting pJMP10397 and pJMP10478 with XhoI and assembling with a piece of synthetic DNA (oJMP1946) encoding an apramycin resistance gene.

### CRISPRtOE Guide Design

Guide RNAs were designed using custom Python scripts (https://github.com/ryandward/CRISPRt_gRNA and https://github.com/ryandward/seq_hunter). The *E. coli* K-12 RefSeq genome assembly, in GenBank file format, was downloaded from the NCBI database (Accession Number: GCF_000005845.2), using the seq_hunter.py script. All genomic locations with a suitable “CN” PAM sequence were identified using the CRISPRt_gRNA.py script. Potential spacers of 32 nucleotides in length were evaluated using awk, and ten were selected based on the following criteria: 1) Unique occurrence in the genome, 2) occurrence on the same strand as the downstream gene, 3) at least 95 nucleotides upstream from the gene annotation.

### CRISPRt/CRISPRtOE individual guide and guide library construction

Guide sequences (Table S5A-C) were cloned into the BsaI-cloning site of the pCRISPRt-T/pCtOE plasmids. For cloning individual spacer sequences, two 36-nucleotide (nt) oligonucleotides (oligos) were annealed so that they encode the desired sequence flanked by sticky ends compatible to the vector BsaI site, whereas for cloning pooled libraries, DNA fragments encoding spacers were amplified from a pool of oligos and digested with BsaI prior to ligation.

To prepare the pCRISPRt-T/pCtOE vectors for cloning, plasmid DNA was extracted from a 100 ml culture using a midiprep kit, 2-10 µg plasmid DNA was digested with BsaI-HF-v2 (NEB R3733) in a 100 µl reaction for 4 h at 37°C and then spin-purified. For cloning individual guides, pairs of oligonucleotides (2 µM each) were mixed in a 50 µl total volume of 1X NEB CutSmart buffer, heated at 95°C, 5 min and then cooled to room temperature ∼20 min to anneal, and then diluted 1:40 in dH_2_O prior to ligation. Ligation reactions (10 µl) contained 1X T4 DNA ligase buffer (NEB), additional DTT (10 mM final) and ATP (0.1 mM final), and 0.5 µl T4 DNA ligase in addition to 50 ng BsaI-digested/spin-purified vector and 2 µl of 1:40 diluted annealed oligos. Ligations were incubated at 25°C for 2 h and ligase was heat inactivated for 20 min at 65°C. 1-2 µl ligation was used for electroporation into 50 µl *E. coli* strain BW25141 (sJMP146). Transformations were serially diluted and plated with selection on carb to obtain isolated colonies. After confirmation of sequence, individual plasmids were transformed into *E. coli* mating strain WM6026 (sJMP424 or sJMP3257) with selection on carb and DAP.

For cloning pooled libraries, fragments were amplified from 90 nt pooled oligos (IDT oPools or Agilent SurePrint pools) using the following conditions per 100 µl reaction (reaction size was adjusted ∼100-600 µl depending on size of library): 20 µl Q5 buffer, 3 µl GC enhancer, 2 µl 10mM each dNTPs, 5 µl each 10 µM forward and reverse primers, 2 µl 10 nM pooled oligo library, 1 µl Q5 DNA polymerase, and 186 µl H_2_O with the following thermocycling parameters: 98°C, 30s; 15-19 cycles of: 98°C, 15s; 56°C, 15s; 72°C, 15s; 72°C, 10 min; 10°C, hold (cycles were adjusted to obtain sufficient PCR product without overamplifying/biasing the library). PCR products (90 bp) were spin-purified, and 300 ng was digested with BsaI-HF-v2 (NEB R3733) in a 100 µl reaction for 2 h at 37°C (no heat kill, reaction size was adjusted proportionally depending on the size of the library). Size and digestion of PCR products were confirmed on a 4% agarose E-Gel (Thermo). The BsaI-digested PCR product without further purification (3ng/µl) was ligated into BsaI-digested, spin-purified plasmid as described above except 10 µl of the digested PCR product was ligated with 500 ng cut plasmid in a 100 µl reaction and ligations were incubated at 16°C for 14 h. Library ligations were purified by spot dialysis on a nitrocellulose filter (Millipore VSWP02500) against 0.1 mM Tris, pH 8 buffer for 20 min prior to transformation by electroporation into *E. coli* mating strain WM6026 (sJMP424 or sJMP3257).

To obtain sufficient transformants for large libraries, electrocompetent cells were made 5X more concentrated in the final step of the preparation(67) and multiple transformations were carried out depending on the size of the library. Transformations were plated with selection on LB with carb and DAP at a density of ∼30,000-50,000 colonies/150 mm petri plate. Cells (>30X more colonies the number of guides, e.g. >1.35 million CFU for the 45,000 guide libraries) were scraped from plates and resuspended in LB + 15% glycerol, density was adjusted to OD_600_=9, and aliquots were frozen at -80°C.

### Transfer of the CRISPRt/CRISPRtOE system to the bacterial chromosome

CRISPRt and CRISPRtOE strains were constructed by triparental mating of two *E. coli* donor strains—one with the pCRISPRt-H plasmid encoding Cas678-TniQ-TnsABC and another with the pCRISPRt-T or pCtOE plasmid containing the transposon with the guide RNA and antibiotic resistance—and a recipient strain. See Table S2 for a summary of growth media and conditions and antibiotic concentrations. All conjugations and selection post-conjugation were carried out at 30°C regardless of the normal incubation temperature for the organism. Isolated donor and recipient strains were struck out on plates with the appropriate media and antibiotic (if relevant) concentrations. Colonies were resuspended from plates into the growth medium used for the recipient to a density of OD_600_=9. Equal amounts of donors and recipient were mixed together (along with the appropriate no recipient, no CRISPRt-H donor, or no CRISPRt-T donor controls), cells were pelleted by centrifugation at 5000 x *g* (except *A. baumannii* at 9000 x *g*), placed on a nitrocellulose filter (Millipore HAWP01300) on an LB plate (except an RMG plate for *Z. mobilis*), and incubated 12-18 h at 30°C. Cells from the filter were resuspended in 300 µl media, serially diluted, and plated with antibiotic selection for the transposon (see growth and selection conditions above) and without DAP to select against the donor strains, and incubated at 30°C until colonies formed (1-3 days). A typical mating was 100 µl each strain and plating 100 µl undiluted, and 10^-1^ through 10^-4^ dilutions to obtain isolated colonies. For library construction, conjugations were scaled up and plated with selection at a density of ∼30,000-50,000 colonies/150 mm petri plate. Cells (>30X more colonies the number of guides, e.g. >1.35 million CFU for the 45,000 guide libraries) were scraped from plates and resuspended in LB + 15% glycerol, density was adjusted to OD_600_=10-15, and aliquots were frozen at -80°C.

### Analysis of phenotype of *E. coli lacZ* and *gfp*-targeting CRISPRt

For *gfp* targeted CRISPRt, fluorescence level (GFP) of 80 individual colonies was determined. Cultures were grown in 300 µl media in 96 well deepwell plates from a single colony to saturation. Cells were centrifuged at 4,000 x *g* for 10 min and cell pellets were resuspended in 300 µl 1X PBS. 150 µl was transferred to a 96-well black, clear bottom microplate (Corning 3631) and cell density (OD_600_) and fluorescence (excitation 475 nm, emission 510 nm) were measured in a fluorescence microplate reader (Tecan Infinite Mplex). Fluorescence values were normalized to cell density. For *lacZ* targeted CRISPRt, the color of 80 individual colonies was determined by patching on LB agar plates containing X-gal (20 µg/ml). Blue indicates the presence of an intact *lacZ* due to metabolism of the X-gal by LacZ (betagalactosidase) and white indicates disruption of *lacZ*.

### Analysis of individual CRISPRtOE isolates from various Proteobacteria

#### Phenotype determination

Fluorescence levels (mScarlet-I) of 12 CRISPRtOE isolates of all 8 species were determined. Cultures of isolates were grown in 300 µl media in 96 well deepwell plates from a single colony to saturation. Cultures were serially diluted to 1:10,000 (1:100 twice) into fresh media and grown again to saturation (∼13 doublings). Cells were centrifuged at 4,000 x *g* for 10 min. Cell pellets were resuspended in 300µl 1X PBS and 150 µl was transferred to a 96-well black, clear bottom microplate (Corning 3631). Cell density (OD600) and fluorescence (excitation 584 nm, emission 607 nms) were measured in a fluorescence microplate reader (Tecan Infinite Mplex). Fluorescence values were normalized to cell density. The assay was repeated three times for *E. coli* and *Z. mobilis* and twice for all other strains.

#### Genotype determination

Insertion positions of 12 CRISPRtOE isolates of all 8 species were determined. Cultures of isolates were grown in 300 µl media in 96 well deepwell plates, serially diluted 1:100 in dH_2_O, heated to 90°C for 3 min prior to use as a template for PCR. Fragments (∼2000 bp) were amplified by PCR with a forward primer in the transposon (oJMP490) and a set of indexed (4 nt barcode) reverse primers in the *mScarlet-I* coding sequence (oJMP2118-2129) in a 25 µl reaction containing: 5 µl OneTaq buffer, 0.5 µl 10mM each dNTPs, 0.5 µl each 10 µM forward and reverse primers, 2 µl diluted culture, 0.25 µl OneTaq DNA polymerase, and 16.25 µl H_2_O with the following touchdown PCR program: 94°C, 3 min then 10 cycles of 94°C, 30s, 65°C, 30s (-1°C/cycle), 68°C, 2.5 min, then 25 cycles of 94°C, 30s, 55°C, 30s, 68°C, 2.5 min, then 68°C, 5 min in a BioRad T100 thermalcycler. Sets of indexed PCRs for each organism were pooled, spin purified and sequenced by Oxford nanopore long read amplicon sequencing by Plasmidsaurus. Sequencing data was demultiplexed using a custom Python script (SplitSamplesSeq.py) and insertion position relative to the *mScarlet-I* ATG start codon was determined.

### mScarlet-I expression analysis

Overnight cultures were diluted 1:1000 into 25ml LB in a 125ml flask in triplicate and incubated at 37°C, shaking until cultures reached mid-log phase (OD600 ∼0.4). Three 3ml culture was treated by vortexing in 375µl cold 5% phenol in ethanol for 10s. Cells were harvested by centrifugation for 10min at 4000xg, flash frozen in a dry ice ethanol bath after decanting supernatant, and stored at -80°C. Cells were lysed by resuspending in 200µl RNase-free TE buffer, pH8 containing 1µl Ready-Lyse lysozyme (Lucigen R1804M) for 5 min at RT and 1min at 55°C. RNA was extracted with the Monarch Total RNA miniprep kit according to the manufacturer’s protocol (includes on column gDNA removal and DNase I treatment). An additional DNA removal step was performed using the DNA-free removal kit (Invitrogen AM1906) according to the manufacturer’s protocol. RNA was quantified using a nanodrop spectrophotometer and Qubit fluorometer (Thermo). cDNA was synthesized using Superscript III Reverse Transcriptase (Invitrogen 18080093) using ∼3µg total RNA and gene-specific primers at 55°C for 30min according to the manufacturer’s protocol. qPCR was performed using a Roche Lightcycler 480 and Lightcycler 480 SYBR Green I Mastermix (Roche 04707516001) in a 20µl reaction containing 10µl 2X mastermix, 1µl each forward and reverse primer (10µM), and 2µl cDNA with preincubation at 95°C 10min and amplification at 95°C 10s, 55°C 15s, 72°C 10s for 40 cycles. Fold changes in mScarlet-I gene expression were calculated using the double delta Ct method (72) with *rho* expression for normalization. Two technical replicates are shown each of with the standard deviation of three biological replicates.

### CRISPRtOE strain construction efficiency

The CRISPRtOE system was transferred to the recipient chromosome by tri-parental conjugation followed by CAST transposition. The reported values are the combined efficiency of conjugation and transposition with the LZ1 guide into a recipient harboring the LP-mScarlet-I reporter. Equal amounts (100µl culture normalized to OD600=9 in media) of recipient, CRISPRt-H donor, and CRISPRt-T donor were mixed and incubated for 16h at 30°C on a nitrocellulose filter on a non-selective plate (RMG for *Z. mobilis*, LB for all other organisms). Cells were resuspended off the filter by vortexing, diluted serially (1:10) in media in triplicate, and 10µl was spotted on selective and non-selective plates and incubated at 30°C. Growth on selective versus non-selective plates was compared to determine efficiency.

### Multiplex CRISPRtOE strain construction

Strains with two CRISPRtOE insertions were constructed using the same conjugation methodology as for single CRISPRtOE insertions except for method 1, two transposon donor strains with different antibiotic resistance cassettes were used. Insertion position of isolates was confirmed by PCR with one primer in the transposon and a second in the target gene and precise location was determined by sequencing the PCR product. Increased antibiotic resistence of the CRISPRtOE isolates was confirmed by disc diffusion assays. MICs of single isolates from each method were determined using MIC test strips (Lsiofilchem).

### CRISPRtOE library growth experiments

#### Competition experiment growth

The *E. coli* CRISPRtOE *murA* and *folA* libraries were mixed in equal volume with the LP-*lacZ* library: 50 µl frozen stock (OD_600_ = 10) of each library (*murA* + LP-*lacZ* and *folA* + LP-*lacZ*) into 100 ml LB (starting OD_600_ = 0.01) in a 500 ml flask and incubated shaking at 37°C until OD_600_ = 0.2 (∼2.5 h) (timepoint = T0) to revive the cells. These cultures were diluted to OD_600_ = 0.02 into 4 ml LB plus antibiotic fosfomycin (0.4 µg/ml, for the *murA* + LP-*lacZ* libraries) or trimethoprim (0.1 µg/ml, for the *folA* + LP-*lacZ* libraries) or no antibiotic control in 14 ml snap cap culture tubes (Corning 352059) in duplicate and incubated with shaking for 18 h at 37°C (T1). These cultures were serially diluted back to OD_600_ = 0.01 into fresh tubes containing the same media and incubated again with shaking for 18 h at 37°C (T2) for a total of ∼10-15 doublings. Cells were pelleted from 15 ml of culture in duplicate at each time point T0 and 1ml of culture at timepoints T1 and T2 and stored at -20°C.

#### Whole genome experiment growth

The *E. coli* CRISPRtOE whole genome libraries with no promoter (sJMP10704) or promoter H (sJMP10705) were revived by dilution of 100 µl frozen stock (OD_600_ = 15) into 100 ml LB (starting OD_600_ = 0.015) and incubation in 500 ml flasks shaking at 37°C until OD_600_ = 0.2 (∼2.5 h) (timepoint = T0). These cultures were diluted to OD_600_ = 0.02 into 4 ml LB plus antibiotic trimethoprim (0.1 µg/ml) or no antibiotic control in 14 ml snap cap culture tubes (Corning 352059) in duplicate and incubated with shaking for 18 h at 37°C (T1). These cultures were serially diluted back to OD_600_ = 0.01 into fresh tubes containing the same media and incubated again with shaking for 18 h at 37°C (T2) for a total of ∼10-15 doublings. Cells were pelleted from 15 ml of culture in duplicate at time point T0 and 1ml of culture at timepoints T1 and T2 and stored at -20°C.

### Analysis of individual CRISPRtOE isolates from library competition experiment

Insertion positions of 8 CRISPRtOE isolates from the *folA* and *murA* libraries were determined. Cultures of isolates were grown in 3 ml media from an isolated colony, serially diluted 1:100 in dH_2_O, heated to 90°C for 3 min prior to use as a template for PCR. Fragments (∼600 bp) were amplified by PCR with a forward primer in the transposon (oJMP61) and a reverse primer in the *folA* or *murA* coding sequence (oJMP2349 and oJMP2348, respectively) in a 50 µl reaction containing: 10 µl OneTaq buffer, 1.0 µl 10mM each dNTPs, 1.0 µl each 10 µM forward and reverse primers, 2 µl diluted culture, 0.5 µl OneTaq DNA polymerase, and 16.25 µl H_2_O with the following touchdown PCR program: 94°C, 3 min then 10 cycles of 94°C, 30s, 65°C, 30s (- 1°C/cycle), 68°C, 1 min, then 25 cycles of 94°C, 30s, 55°C, 30s, 68°C, 1 min, then 68°C, 5 min in a BioRad T100 thermalcycler. PCR products were spin-purified and Sanger sequenced to determine the transposon insertion position relative to the *mScarlet-I* ATG start codon.

### Tnseq analysis of CRISPRt/CRISPRtOE constructs

The TnSeq protocol was adapted from Klompe et al(31). Genomic DNA (gDNA) was extracted from the equivalent of 1 ml of cells at OD_600_=3 (∼2×10^9^ cells) either resuspended from a plate or from liquid culture and was further purified by spot dialysis on a nitrocellulose filter (Millipore VSWP02500) against 0.1 mM Tris, pH 8 buffer for 20 min. 1 µg gDNA was digested with 4U MmeI (NEB) in a 50 µl reaction at 37°C, 12 hrs and heat inactivated at 65°C, 20 min. The digest was purified using 1.8X magnetic beads (Omega) following the manufacturer’s protocol, eluting in 20 µl 10 mM Tris, pH 8.0. Adapter oligos oJMP1995 and P-oJMP1996 (phosphorylated) were annealed by mixing equal volume of 100µM oligos in 10mM Tris, pH8 (50 µM each), heating to 95°C, 5 min followed by cooling to RT ∼15 min, and diluted 1:10 (5 µM each) for use. Annealed oligos were ligated onto MmeI-digested DNA in a 20 µl reaction with 2 µl 10X T4 ligase buffer, 2 µl 100 mM DTT, 2 µl 1 mM ATP, 12 µl purified MmeI-digested gDNA (50 ng/µl, 600 ng total), 1 µl 5 µM annealed adapter oligos (250 nM final) and 1 µl T4 DNA ligase. Ligations were incubated 14 h at 16°C and the enzyme was heat-inactivated for 20 min at 65°C. Ligations were purified using 1.8X magnetic beads, eluting in 22 µl 10 mM Tris, pH 8. Fragments were amplified by PCR with primers containing partial adapters for index PCR with Illumina TruSeq adapters in a 100µl reaction containing: 20 µl 5X Q5 buffer, 3 µl GC enhancer, 2 µl 10 mM each dNTPs, 5 µl each forward primer (oJMP1997) and reverse primer (oJMP1998 or indexed primers oJMP2022-2033), 10 µl purified ligation, 1 µl Q5 DNA polymerase, and 54 µl dH_2_O with the following program: 98°C, 30s then 18-20 cycles of 98°C, 15s, 66°C, 15s, 72°C, 15s, then 72°C, 10 min in a BioRad C1000 thermal cycler. PCR products were spin-purifed (eluted in 15 µl) and quantified fluorometrically. Samples were sequenced by the UWBC NGS Core facility or Azenta Amplicon-EZ service. Briefly, PCR products were amplified with nested primers containing i5 and i7 indexes and Illumina TruSeq adapters followed by bead cleanup, quantification, pooling and running on a NovaSeq X Plus (150 bp paired end reads) or MiSeq (250 bp paired end reads). Sequencing analysis of the initial CRISPRt or CRISPRtOE plasmid libraries from which the strain libraries were prepared was by amplification with oJMP2011 and oJMP1998 (or barcoded version of oJMP1998: oJMP2022-2033), followed by spin purification and Illumina sequencing.

### Analysis of *E. coli mScarlet-I*-targeting CRISPRt libraries

Two plasmid libraries (pJMP10505 and pJMP10506) were constructed (gen^R^ and kan^R^) and used to create CRISPRtOE libraries (sJMP10519, 10520, 10594, 10595, and 10637-10640) in each of four *E. coli mScarlet-I* reporter strains (sJMP10205, 10269, 10630, 10633). Plasmid DNA from pJMP10505 and pJMP10506 was amplified with oJMP2011 and oJMP1998. gDNA from (sJMP10519, 10520, 10594, 10595, and 10637-10640) was processed by TnSeq as detailed above with oJMP1997 and oJMP2022-2033. Sequencing data was demultiplexed using SplitSamplesSeq.py and guides were quantified using seal.sh from bbmap.

### MIC assays

The minimal inhibitory concentration (MIC) of *E. coli folA* and *murA* CRISPRtOE isolates was determined by either growth in a microtiter plate or by disc diffusion assays, respectively. For the broth microdilution assay, CRISPRtOE isolates and WT controls were grown to saturation from an isolated colony and serially diluted to OD600 = 0.003) and the CRISPRtOE library was diluted from a glycerol stock to the same density. Trimethoprim was serially diluted in DMSO at 1000X concentration and then diluted to 2X in MHB. *E. coli folA* CRISPRtOE isolates, the *folA* CRISPRtOE library, a *lacZ* CRISPRtOE control, and WT cultures and the media containing antibiotic were mixed in equal proportions and incubated 20 h shaking at 37°C prior to cell density (OD_600_) determination in a microplate reader. For the MIC strip assays, *E. coli folA* or *murA* CRISPRtOE isolates, a *lacZ* CRISPRtOE control, and WT cultures were diluted to OD_600_ = 0.3 and 300 µl was spread on a 100 mm MH agar plate and dried in a laminar flow cabinet. When dry, MIC strips (Liofilchem, Italy; 920371 (TMP 0.002-32 mg/L) and 920791(FOS 0.064-1024 mg/L)) were applied to the plate and 10 µl serially diluted antibiotics were applied to the discs. Plates were incubated at 37°C for 18 h. Broth microdilution assays were repeated 3 times and MIC test strip assays twice.

### CRISPRtOE data processing and analysis

All commands and scripts used for processing and analysis of CRISPRtOE data are available at (https://github.com/GLBRC/CtOE_analysis).

#### Paired-end analysis for spacer-insertion site analysis

FASTQ files were separated for each read (R1 and R2) and trimmed to remove transposon-specific sequences using Cutadapt (v5.0)(73). The ‘--discard-untrimmed’ flag was used to remove reads that lacked transposon-specific adapters. R1 reads were filtered to retain only trimmed sequences of length 18-20bp, and R2 reads were filtered to retain only those of length 32bp. Reads were then repaired using BBMap (v39.06) suite’s ‘repair.sh’ tool with options to discard unpaired reads (‘repair=t tossbrokenreads=t’). This ensured that only properly paired reads with both R1 and R2 passing all quality filters were retained for downstream analysis. The trimmed FASTQ files were separately aligned to the *E. coli* genome (NCBI RefSeq Assembly ID GCF_000005845.2) or the *Z. mobilis* genome (NCBI RefSeq Assembly ID GCF_003054575.1) using Bowtie (v1.3.1) (74) in single-end mode, with flags ‘-k 1’, ‘--best’, ‘--stratà, ‘--nomaqround’, ‘-v 0’, and ‘-m 1’ . The alignment SAM file for each sample was converted to BAM using Samtools (v1.13)(75), filtered to retain alignments with MAPQ of >=42 with ‘samtools view’, then namesorted with ‘samtools sort –n’, before being parsed to TSV format for downstream analysis. (76) A custom python script was used to annotate the site of Tn insertion, identify unique sites of Tn insertion, and compare these to the predicted sites of Tn insertion from the designed CRISPRtOE library. <u>CRISPRtOE library sequencing analysis (single-end)</u>. Prior to analysis of FASTQ reads, a list of expected Tn insertion sites based on the CRISPRtOE library design was created with a custom python script (ctoe_analysis ‘guides’ pipeline). R1 reads were trimmed and filtered with Cutadapt as in paired-end analysis. Trimmed R1 reads were aligned to reference genome with bowtie, then filtered and sorted with SAMtools as in paired-end analysis. Alignment files were parsed to TSV format, sites of Tn insertion identified and counted with a custom python script. Observed sites of Tn insertion were compared to predicted sites of Tn insertion. Counts of insertions consistent with the target location and orientation of CRISPRtOE library design were retained for downstream analysis in edgeR.

### Differential abundance analysis

Differential abundance analysis was performed using the generalized linear model (GLM) framework in edgeR (v4.4.2). Raw count data were filtered to retain unique Tn insertions with counts per million (CPM) of at least 0.1 in a minimum of half of the samples per comparison set. Normalization factors were calculated using the trimmed mean of M-values method. A negative binomial GLM was fitted using quasi-likelihood (QL) methods with robust dispersion estimation. The model incorporated both treatment and library type factors, modeling each independently as well as their interaction (formula: ∼ library * treatment). Differential abundance was assessed for individual insertions. For gene-level analysis, a competitive gene set test was performed using CAMERA, linking Tn insertions to their downstream genes. Separate contrasts were constructed to evaluate treatment effects in promoter and non-promoter contexts, as well as the interaction between library type and treatment. For each gene, median log fold changes were calculated from the insertions mapping to that gene. Multiple testing correction was performed using the Benjamini-Hochberg procedure to control the false discovery rate. Downstream gene-level analyses report the median log fold change calculated from individual insertions and the FDR values calculated using CAMERA. Genes with significant log2FCs between treatment and control conditions were identified from CAMERA analysis, requiring an FDR <= 0.05. Genes with promoter-dependent phenotypes were selected, requiring the median log2FC for the library * treatment interaction factor to be at least 2 (either positive or negative). Genes were then ranked by the log2FC value of the Promoter library, and the top 10 and bottom 10 genes selected for plotting.

## Data availability

Raw sequencing data is deposited at the Sequence Read Archive (https://www.ncbi.nlm.nih.gov/sra, PRJNA1284861). All other data are available upon request. Plasmids are available from Addgene (https://www.addgene.org, #225832-225871). Guide design script is available at https://github.com/ryandward/CRISPRt_gRNA. Analysis and demultiplexing scripts are available at https://github.com/GLBRC/CtOE_analysis and https://github.com/GLBRC/CtOE_downstream_analysis. (Pre-publication SRA reviewer link: https://dataview.ncbi.nlm.nih.gov/object/PRJNA1284861?reviewer=8p8ge077d7g4jthl52a6tuadmv)

## Supplemental Material

Supplemental Figures S1-S17

Supplemental Tables S1-S12

## Acknowledgements

We thank members of the Peters lab and Michael Thomas for helpful discussions, Joshua Hyman and the University of Wisconsin-Madison Biotechnology Center for technical assistance with Illumina sequencing, and Samuel Sternberg for plasmids. This material is based upon work supported by the Great Lakes Bioenergy Research Center, U.S. Department of Energy, Office of Science, Biological and Environmental Research Program under Award Number DE-SC0018409. This work was also supported by the National Institutes of Health 1R35GM150487-01 to J.M.P. and by the Advanced Research Projects Agency for Health, under award number 1AYSAX000005-01 to J.M.P.

## Author contributions

A.B.B. and J.M.P. designed, performed, and interpreted the experiments and wrote the manuscript with input from all authors. K.S.M. and A.N.H. developed the data analysis pipeline and performed data analysis. R.D.W. and M.P. developed code. B.C.D, R.A.C., C.C.F., and E.E.B performed experiments.

---

**Figure S1.**
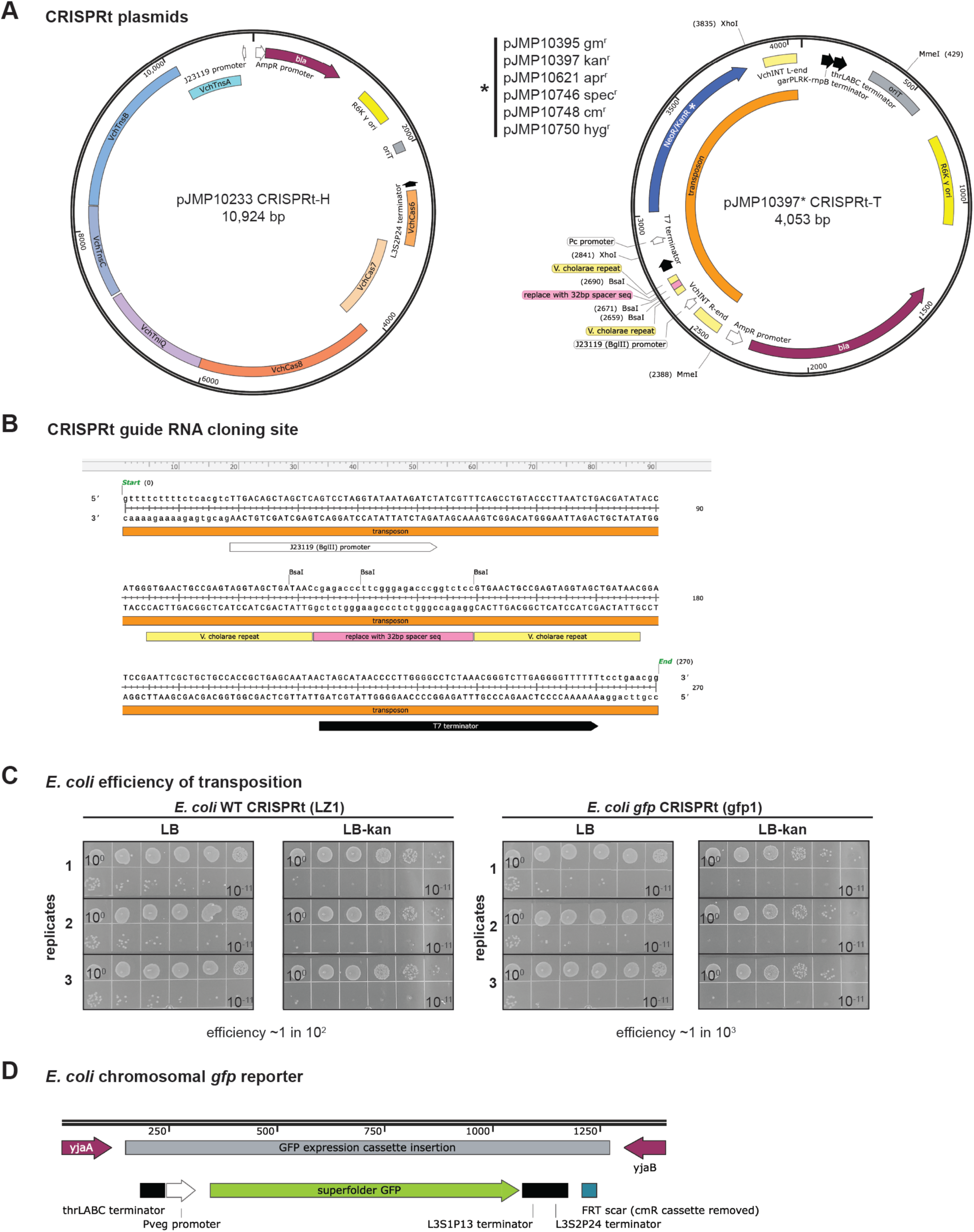
Details of CRISPRt plasmids and *E. coli* GFP reporter strain. (A) Plasmid maps of CRISPRt-H helper and CRISPRt-T transposon plasmids. Both plasmids in the dual plasmid system have backbones with an R6K pir-dependent replication origin, an ampicillan resistance cassette, and conjugative transfer origin. The CRISPRt-H helper plasmid has expression cassettes for the minimal Vch CRISPR (Cas678) and Transposase (TnsABC, TniQ) machinery derived from pSpin-R(21). The CRISPRt-T transposon plasmid encodes a modified VchINT transposon derived from pSpin-R(21) which includes a guide RNA expression cassette with a modified BsaI cloning site and an antibiotic expression cassette and transcription terminators derived from CRISPRi plasmids(71). The antibiotic resistance cassette is flanked by XhoI sites to facilitate modification, six variants are shown. (B) Sequence of CRISPRt targeted transposition system guide RNA expression cassette including the BsaI cloning site for new guide RNAs. (C) Efficiency of obtaining CRISPRt transconjugants in *E. coli* using the *LZ1* and *gfp1* guides. Serial dilutions of three technical replicates of *LZ1* and *gfp1* CRISPRt assays were plated with and without selection (LB-kan or LB, respectively) and CFUs were compared; Efficiency of CRISPRt targeted transposition in *E. coli* with the *LZ1* guide was ∼1 in 100 and with the *gfp1* guide was ∼1 in 1000. (D) Position of the P_veg_-*sfgfp* expression cassette inserted between *yjaA* and *yjaB* on the *E. coli* chromosome.

**Figure S2.**
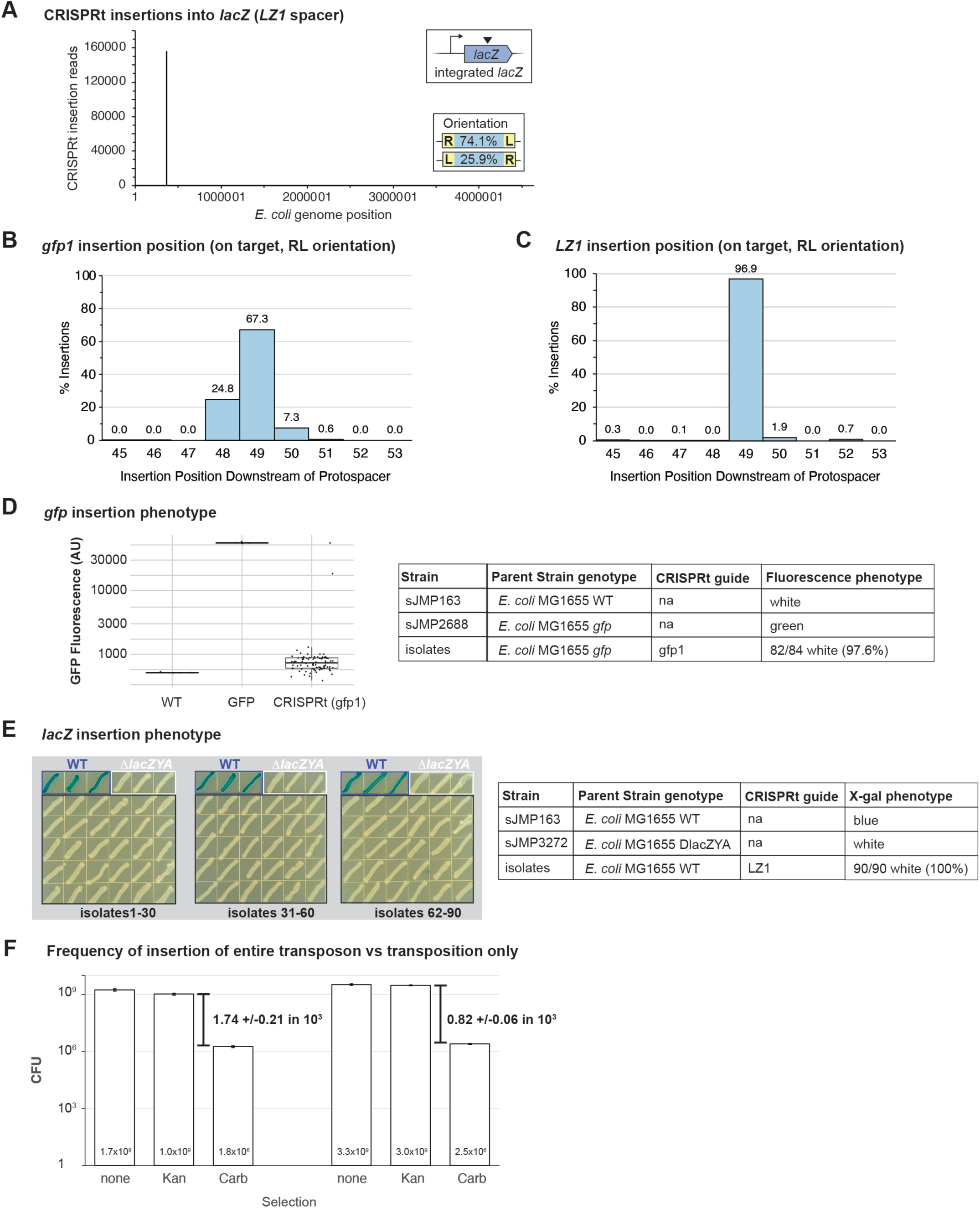
CRISPRt disruption efficiency in *E. coli*. (A) Specificity of CRISPRt disruption of *lacZ* in *E. coli* measured by Tnseq. Mapping of location of transposon insertion sites to the *E. coli* genome after CRISPRt targeted transposition the *LZ1* guide (216,801 reads). (B) Insertion position (bp downstream of protospacer) frequency of CRISPRt disruption of *gfp* and (C) *lacZ* in *E. coli* (RL orientation). Percentage of on-target, RL orientation insertions for all positions with > 0.01%. (D) Efficiency of CRISPRt disruption of *sfgfp* in *E. coli* using a fluorescence phenotypic assay. The *E. coli yjaA*-*sfgfp*-*yjaB* strain (sJMP2688) is green fluorescent whereas the *E. coli* WT strain (sJMP163) is not. Isolates from CRISPRt targeted transposition of *E. coli yjaA*-*sfgfp-yjaB* with the *gfp1* guide (selection for transconjugants on LB-kan plates) were assayed for fluorescence; 82/84 (97.6%) were white indicating disruption of *sfgfp*. (E) Efficiency of CRISPRt disruption of *lacZ* in *E. coli* using an X-gal phenotypic assay. The *E. coli* WT strain (sJMP163) can metabolize X-gal resulting in a blue colony color whereas the *E. coli lacZYA* deletion strain (sJMP3272) cannot metabolize X-gal resulting in a white colony color. Isolates from CRISPRt targeted transposition of *E. coli* WT with the *LZ1* guide (selection for transconjugants on LB-kan plates) were assayed for colony color; 90/90 (100%) were white indicating disruption of *lacZ*. (F) Frequency of co-integration of transposon-containing plasmid versus transposition of transposon only. *E. coli* CRISPRtOE libraries (sJMP10705 and sJMP10706) were serially diluted and plated on LB, LB-kan, and LB-carb in triplicate to determine total CFU, CFU with selection for transposon, CFU with selection for insertion of entire transposon-containing plasmid.

**Figure S3.**
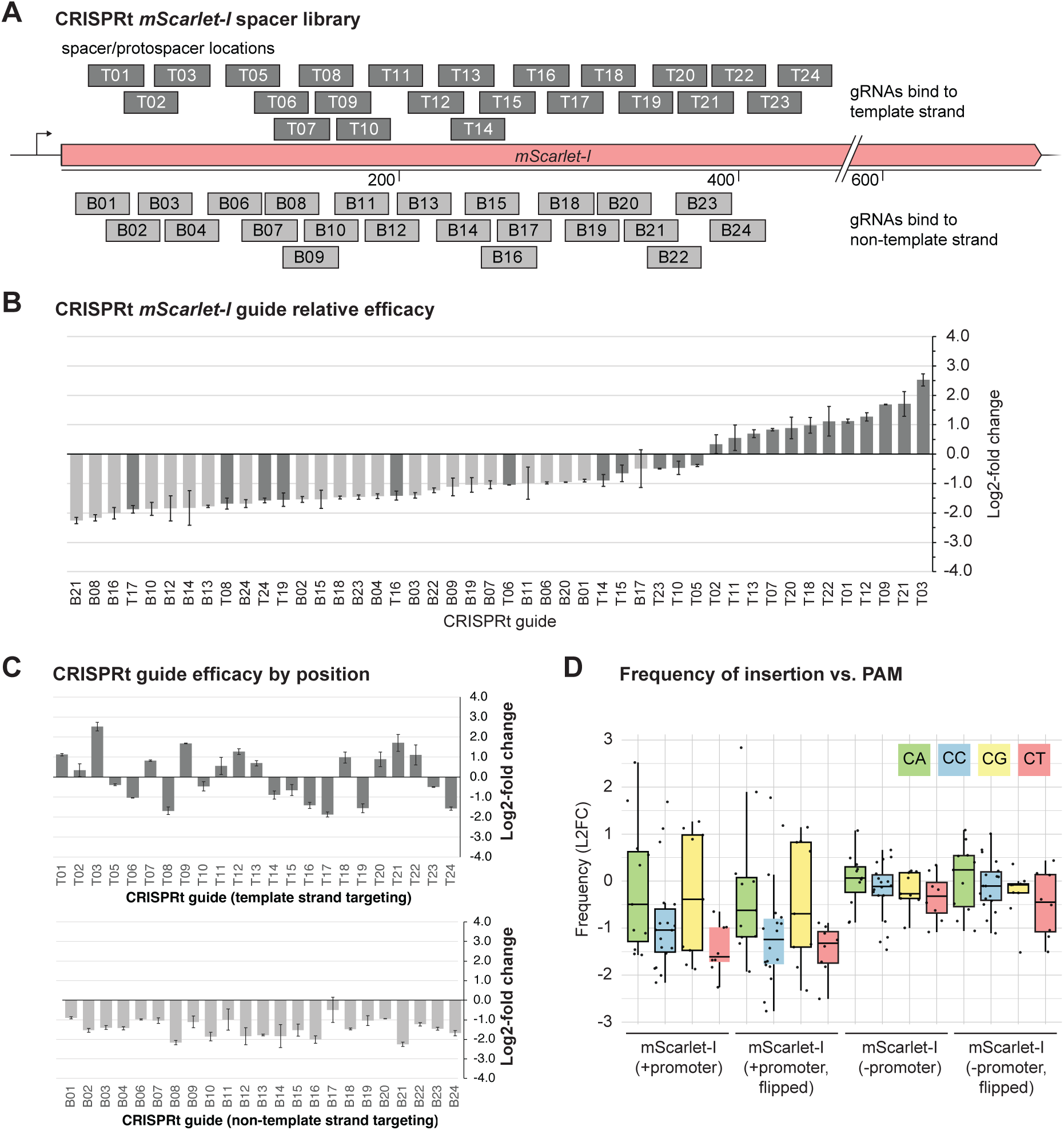
Transcription impacts efficacy of CRISPRt guides. (A) Schematic of guides in pooled *mScarlet-I* disruption experiment. Twenty-three guides matching protospacers located on top (T1-24) or bottom (B1-24) strand of the mScarlet-I encoding gene with a range of CN guides (see Table S5 for details). (B) Relative efficacy of CRISPRt *mScarlet-I* guides in a pooled screen targeting *mScarlet-I* expressed from a transposon in the *att*_Tn*7*_ site on the *E. coli* chromosome. Relative efficacy is expressed as the log_2_-fold change of the frequency in the CRISPRt libraries (sJMP10519 and sJMP10520) vs. the frequency in the original plasmid construct libraries (sJMP10505 and sJMP10506). Average and standard deviation of two separate CRISPRt libraries (gentamicin^R^ and kanamycin^R^) containing all 46 guides. (C) Relative efficacy data ordered by guide position rather than efficacy shows no correlation between position and efficacy. (D) No correlation between PAM sequence (CA, CC, CG, CT) and efficacy (frequency of insertion) for 46 *mScarlet-I* targeting guides.

**Figure S4.**
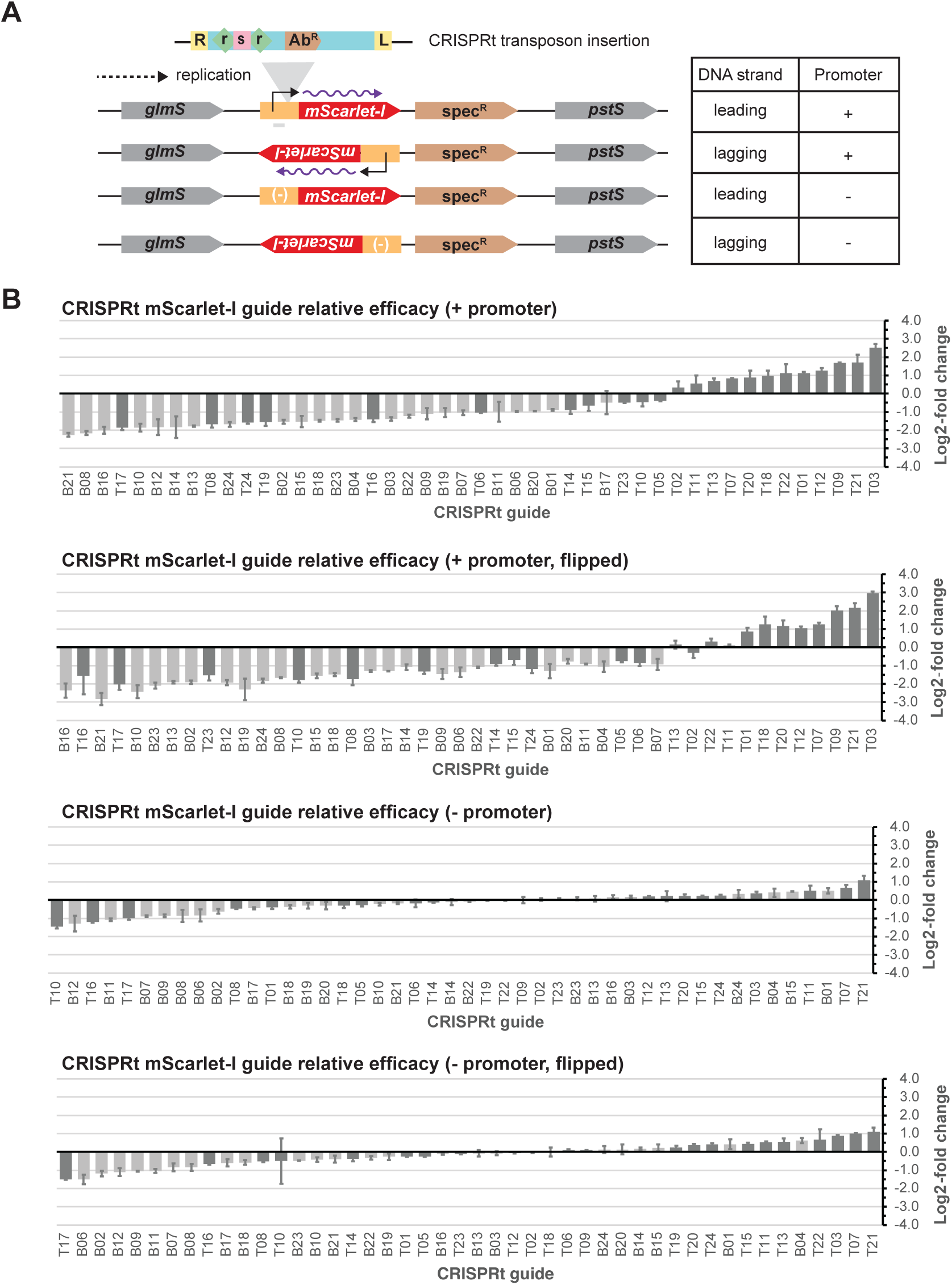
Transcription impacts efficacy of CRISPRt guides. (A) Schematic of *mScarlet-I* cassettes +/- promoter on the *E. coli* chromosome. The *mScarlet-I* expression cassette (red) with and without a promoter (orange) along with a spectinomycin resistance cassette (spt^R^, brown) are located on the *E. coli* chromosome just downstream of *glmS* and upstream of *pstS*. The *mScarlet-I* expression cassette is inserted in one of two orientations so that the direction of transcription is either in the same or the opposite direction of the leading strand of chromosomal DNA replication. (B) Relative efficacy of CRISPRt *mScarlet-I* guides in a pooled screen targeting *mScarlet-I* expressed on the *E. coli* chromosome in different orientations with and without transcription. Relative efficacy is expressed as the log2-fold change of the frequency in the CRISPRt libraries (sJMP10519-10520, sJMP10594-10595, sJMP10637-10640) vs. the frequency in the original plasmid construct libraries (sJMP10505 and sJMP10506) with average and standard deviation of two separate CRISPRt libraries (gentamicin^R^ and kanamycin^R^) containing all 46 guides shown.

**Figure S5.**
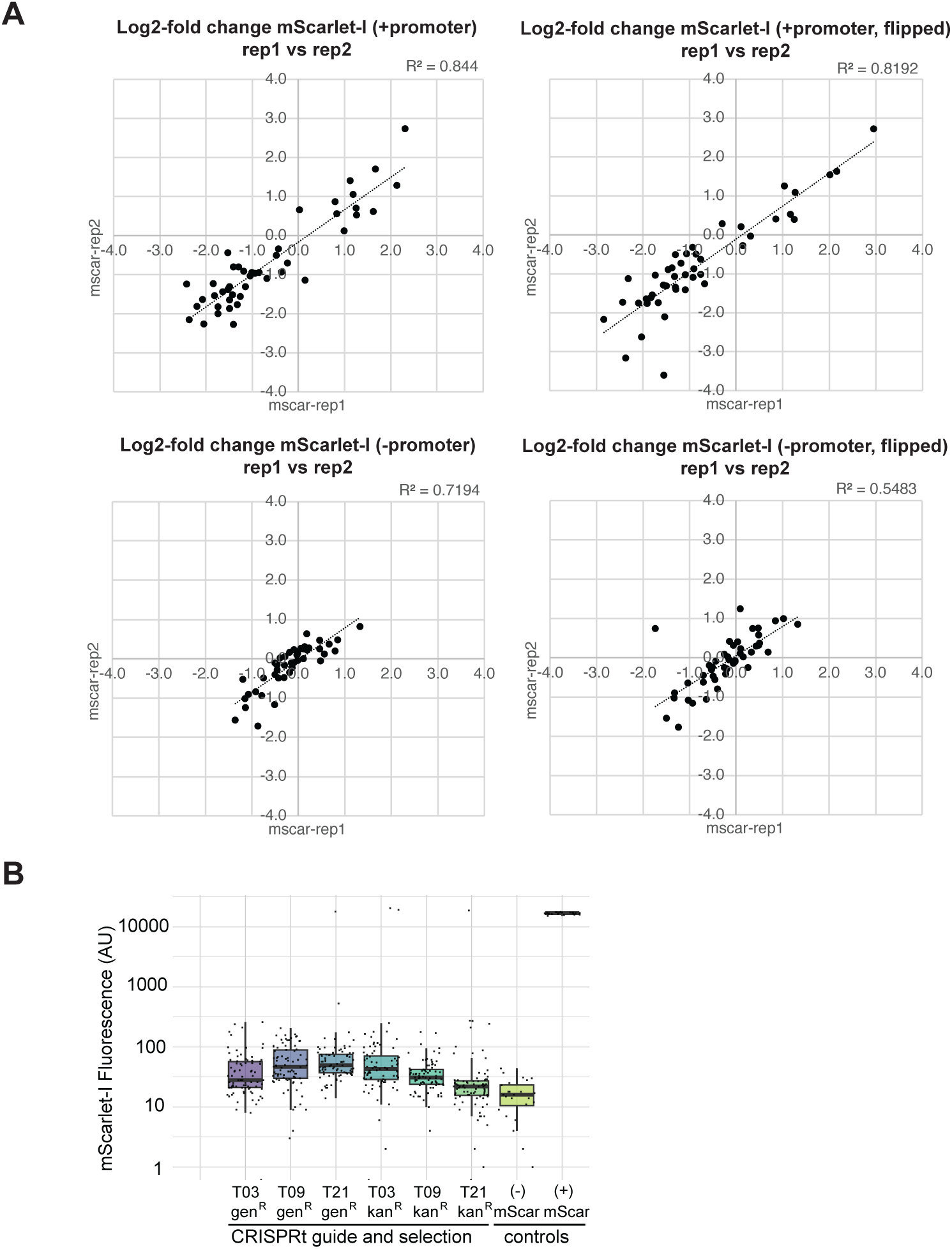
Transcription impacts efficacy of CRISPRt guides, continued. (A) Relative efficacy of CRISPRt gRNAs in a pooled screen targeting *mScarlet-I* as in Figure S4 with average and standard deviation of two separate CRISPRt libraries (gentamicin^R^ and kanamycin^R^) containing all 46 guides shown on scatter plots. (B) Fluorescent phenotypes of 80 isolates from each of the *E. coli* CRISPRt *mScarlet-I* individual guide libraries (sJMP10641-10646) compared to *E. coli* strains with no *mScarlet-I* gene ((-) mScar, sJMP6999) or expressing *mScarlet-I* ((+) mScar, sJMP10205). Only 4/480 (0.83%) retained *mScarlet-I* levels equal to the parent strain indicating an off-target rate of <1% overall.

**Figure S6.**
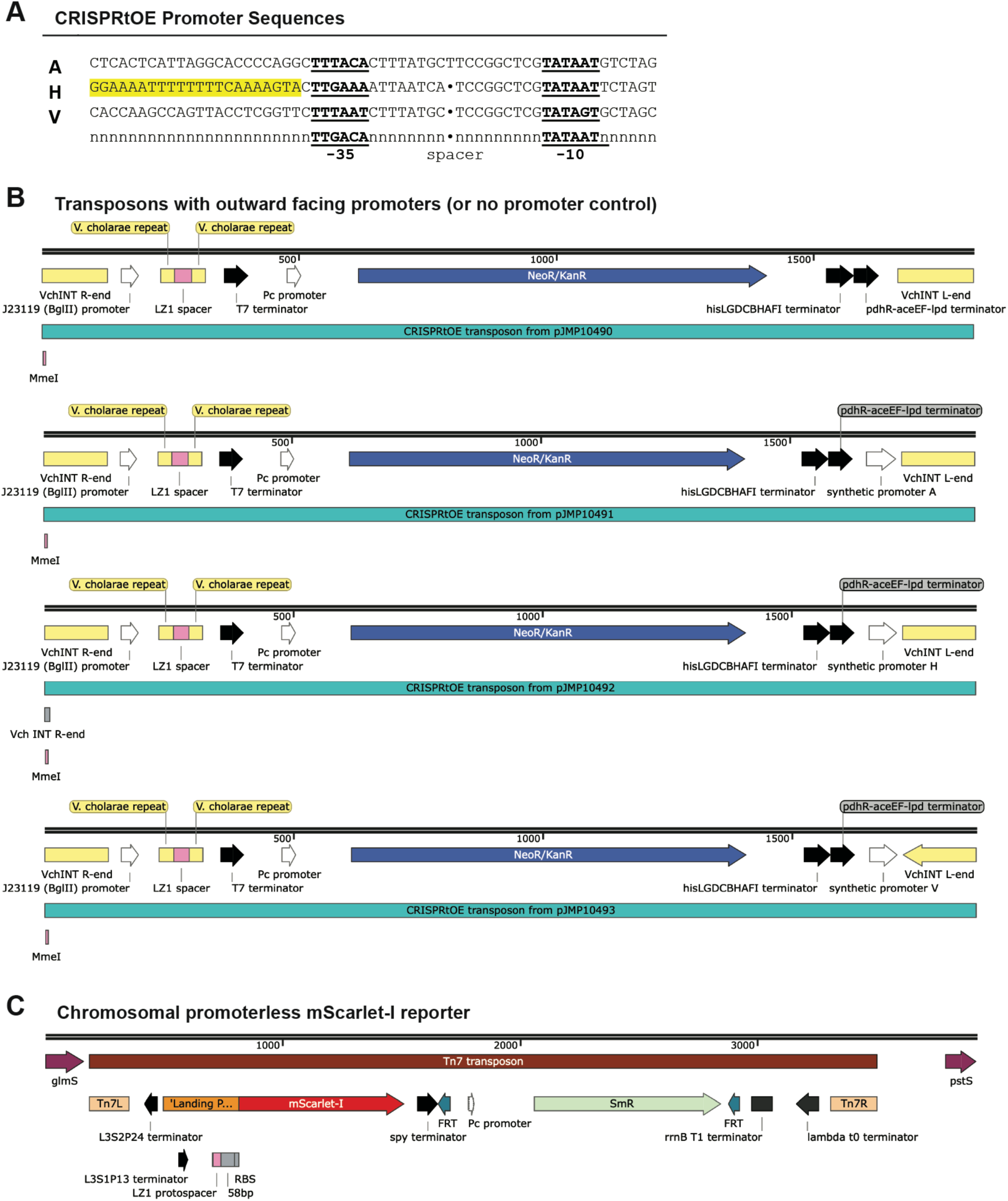
CRISPRtOE construct details. (A) Sequence alignment of CRISPRtOE promoters A, H, and V compared to a consensus *E. coli* σ^70^ promoter. Promoter A is based on the *lacUV5* promoter. Promoter H and V are synthetic promoters. Promoter H has an UP element highlighted in yellow. (B) Schematics of CRISPRtOE Tn*7*-like transposon constructs with four different outward facing promoter sequences (no promoter, and constitutive promoters A, H, and V; see Table S3 for plasmid details). Plasmids have been submitted to Addgene (ID 225832-225871). (C) Schematic of the chromosomally-encoded promoterless *mScarlet-I* reporter contained on a Tn*7* transposon inserted into the *att*_Tn*7*_ site of *E. coli*.

**Figure S7.**
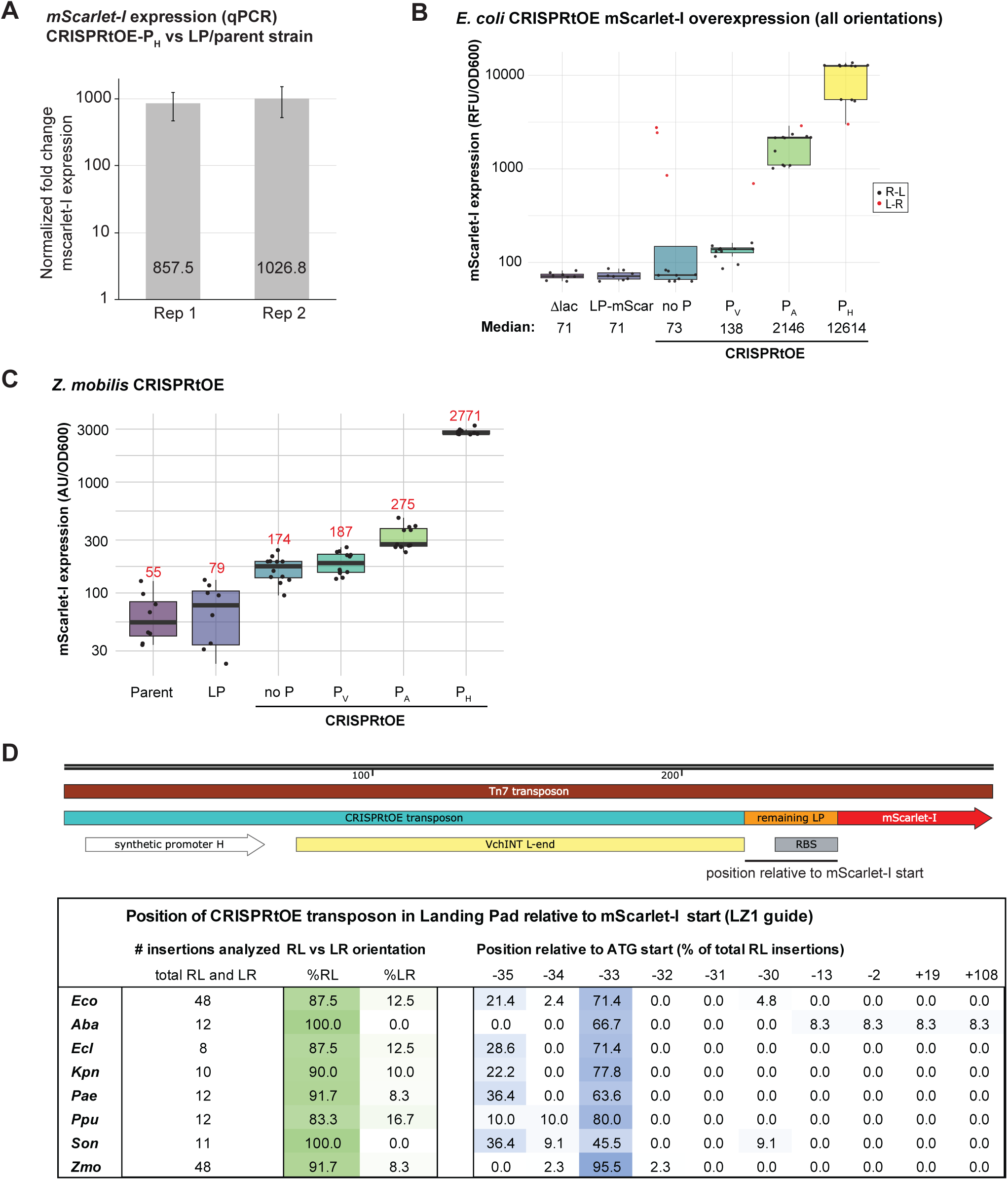
Tunable overexpression of chromosomally located genes using CRISPRtOE. (A) Quantification by qPCR of the *E. coli* CRISPRtOE-P_H_-*mScarlet-I* strain versus the promoterless ’Landing Pad’ parent strain. Assay was repeated twice each with three biological replicates. *mScarlet-I* expression was normalized to expression of *rho*. (B) mScarlet-I fluorescence analysis of *E. coli* CRISPRtOE isolates (no promoter or synthetic promoters A, V, and H) compared to the parent (promoterless *mScarlet-I*) strain. As in Figure 2B except values are shown for all CRISPRtOE isolates (both RL and LR insertion orientations with RL shown in black and LR (flipped) shown in red). Error is expressed for the median value of 12 isolates in three replicate assays. (C) mScarlet-I fluorescence analysis of CRISPRtOE isolates of the Alphaproteobacterium *Zymomonas mobilis* (no promoter or synthetic promoters A, V, and H) compared to the parent (promoterless *mScarlet-I*) strain. Values are shown for isolates with on-target, RL orientation CRISPRtOE insertions. Error is expressed for the median value of 8-12 isolates in three replicate assays. (D) Schematic showing insertion of CRISPRtOE transposon into the ‘Landing Pad’ upstream of *mScarlet-I*. Insertion position is variable (∼45-55bp downstream of protospacer) resulting in a variable amount of the Landing Pad (containing the RBS) remaining upstream of the *mScarlet-I* coding sequence. Insertion positions and orientation of CRISPRtOE isolates in the eight species shown in Figure 2C. Insertion position is expressed relative to start of *mScarlet-I* coding sequence (equal to the remaining LP sequence).

**Figure S8.**
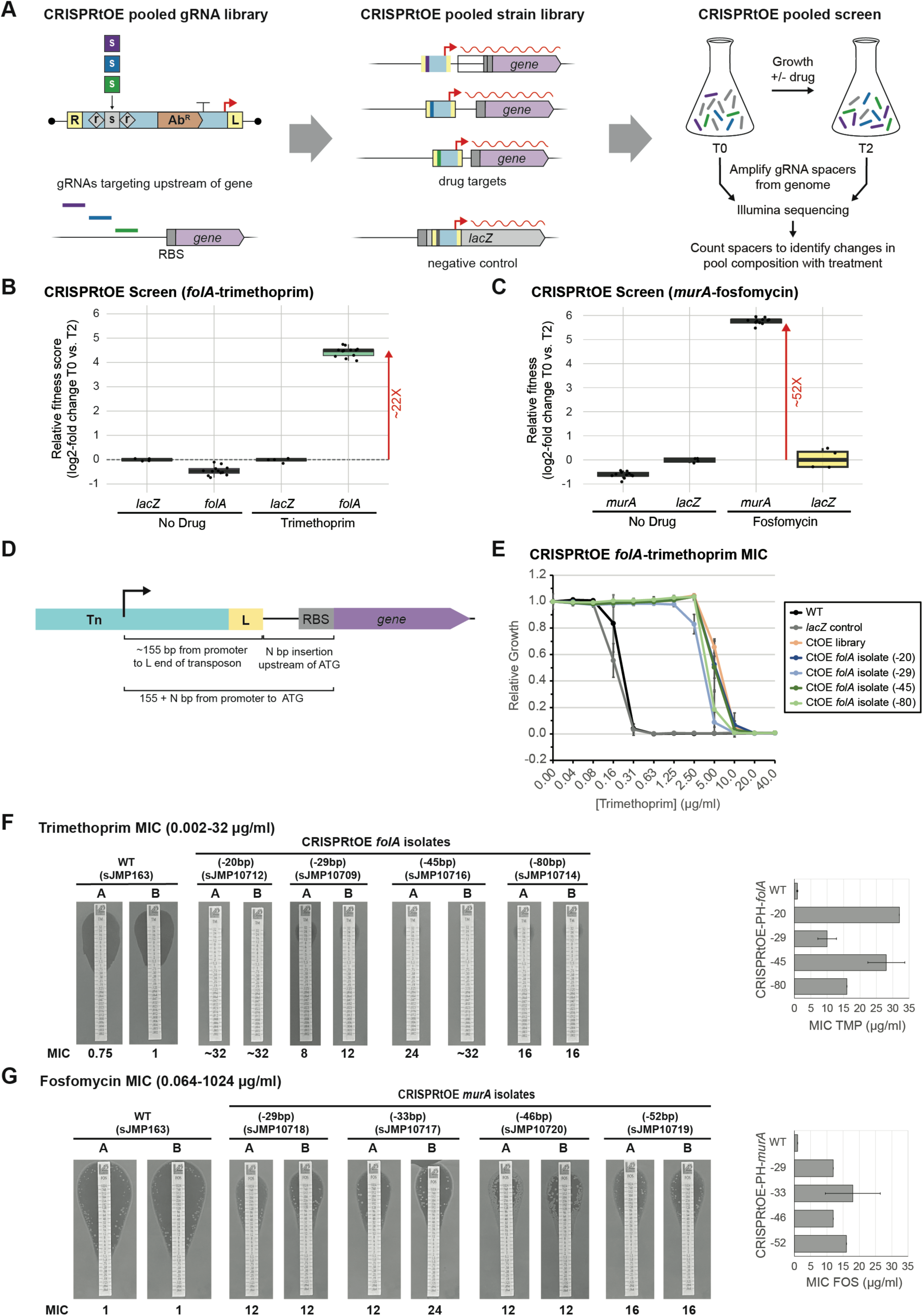
CRISPRtOE *folA* and *murA* pooled library screen and MIC assays inform antibiotic mode of action. (A) Schematic of CRISPRtOE competitive growth assay. A pooled CRISPRtOE strain library was constructed by amplification of spacer sequences from a pooled oligo library and transfer onto the chromosome of a recipient strain. Insertion of the CRISPRtOE transposon is either upstream of genes of interest (12 guides/gene) or within the *lacZ* gene as a control gene (4 guides/gene) in *E. coli*. A CRISPRtOE pooled library was screened by culturing a CRISPRtOE pooled library and control library together for multiple generations in the presence or absence of a chemical of interest (trimethoprim (TMP) for the *folA* library and fosfomycin (FOS) for the *murA* library). The change in the composition of the pooled library before and after treatment was measured by amplification and NGS sequencing of the spacer sequences. (B) CRISPRtOE *folA* pooled library screen in TMP. Fitness of pooled strains overexpressing *folA* compared to the *lacZ* control strains in the absence or presence of TMP. (C) CRISPRtOE *murA* pooled library screen in FOS. (D) Schematic of insertion position for CRISPRtOE isolates. (E) MIC broth microdilution assay of four individual CRISPRtOE *folA* strains compared to WT, a CRISPRtOE *lacZ* isolate, and the CRISPRtOE *folA* pooled library in the presence of TMP (0.04-20 µg/ml). Error is expressed for 3 separate assays. (F) MIC test strip assays (trimethoprim, 0.002-32 µg/ml) comparing WT and four distinct CRISPRtOE *folA* isolates quantified as average and standard deviation of two assays. (G) MIC test strip assays (fosfomycin, 0.064-1024 µg/ml) comparing WT and four distinct CRISPRtOE *murA* isolates quantified as average and standard deviation of two assays.

**Figure S9.**
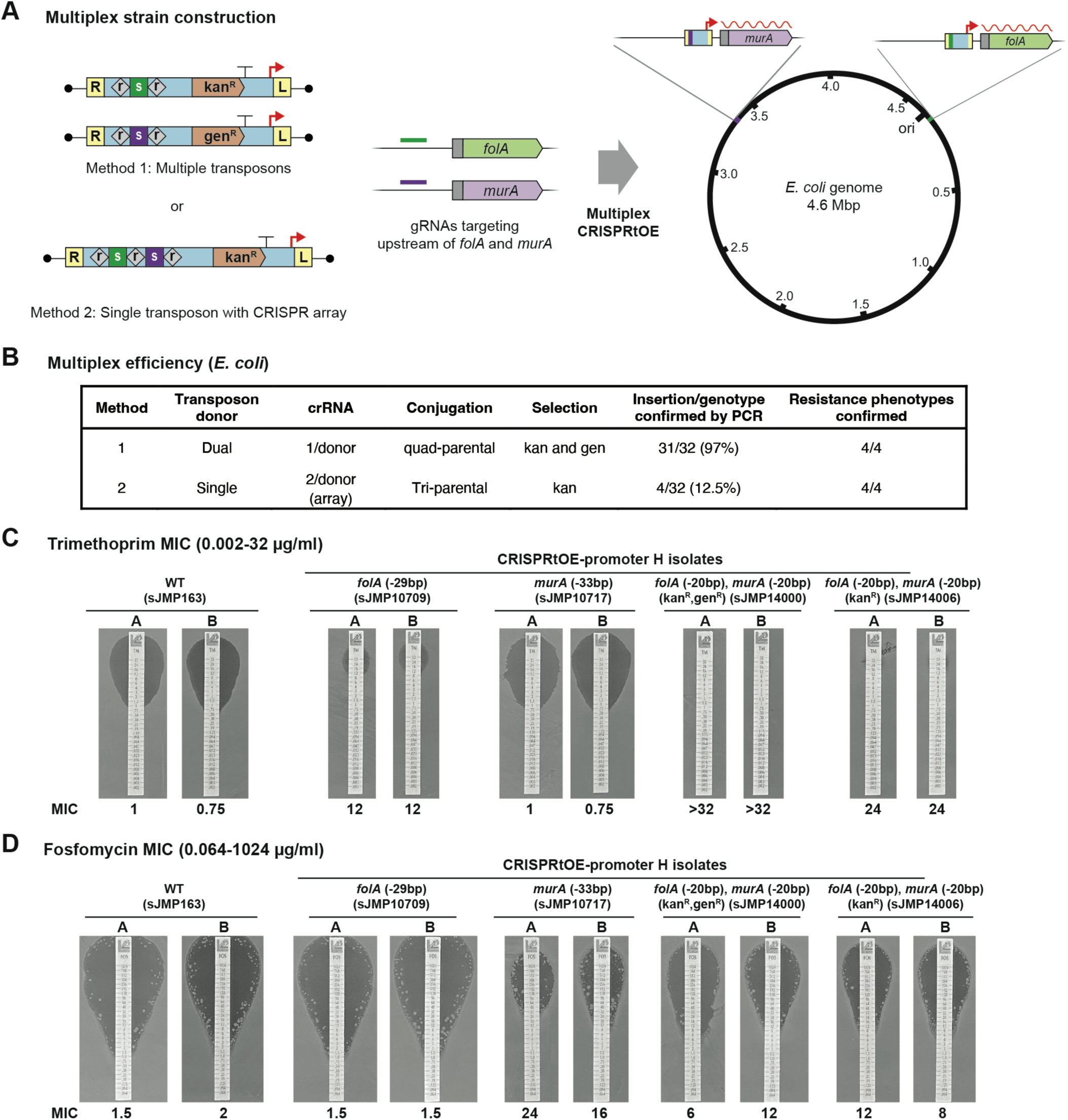
Multiplexed CRISPRtOE strain construction. (A) Schematic of two methods of multiplexed CRISPRtOE strain construction in *E. coli*. Method 1 utilizes quad-parental mating with two CRISPRtOE transposon donors with different antibiotic resistance genes while Method 2 utilizes tri-parental mating with a single transposon donor with a single antibiotic resistance gene and a CRISPR array encoding multiple guides. Guides upstream of *folA* or *murA* that were previously shown to be effective at conferring TMP or FOS resistance (see Figure S8) were used to simultaneously guide the CRISPRtOE-promoter H Tn insertion into the genome of *E. coli*. (B) Efficiency of multiplexed insertion of CRISPRtOE-promoter H-*folA* and CRISPRtOE-promoter H-*murA* as determined by PCR of 32 isolates. Increased TMP and FOS resistance of 4 PCR-confirmed isolates was confirmed by disc diffusion assays. (C) TMP resistance of 1 sequence-confirmed isolate from each of the two methods was compared to WT and previously isolated single isolates (either CRISPRtOE-promoter H-*folA* or CRISPRtOE-promoter H-*murA*. (D) FOS resistance of 1 sequence-confirmed isolate from each of the two methods was compared to WT and previously obtained single isolates (either CRISPRtOE-promoter H-*folA* or CRISPRtOE-promoter H-*murA*.

**Figure S10.**
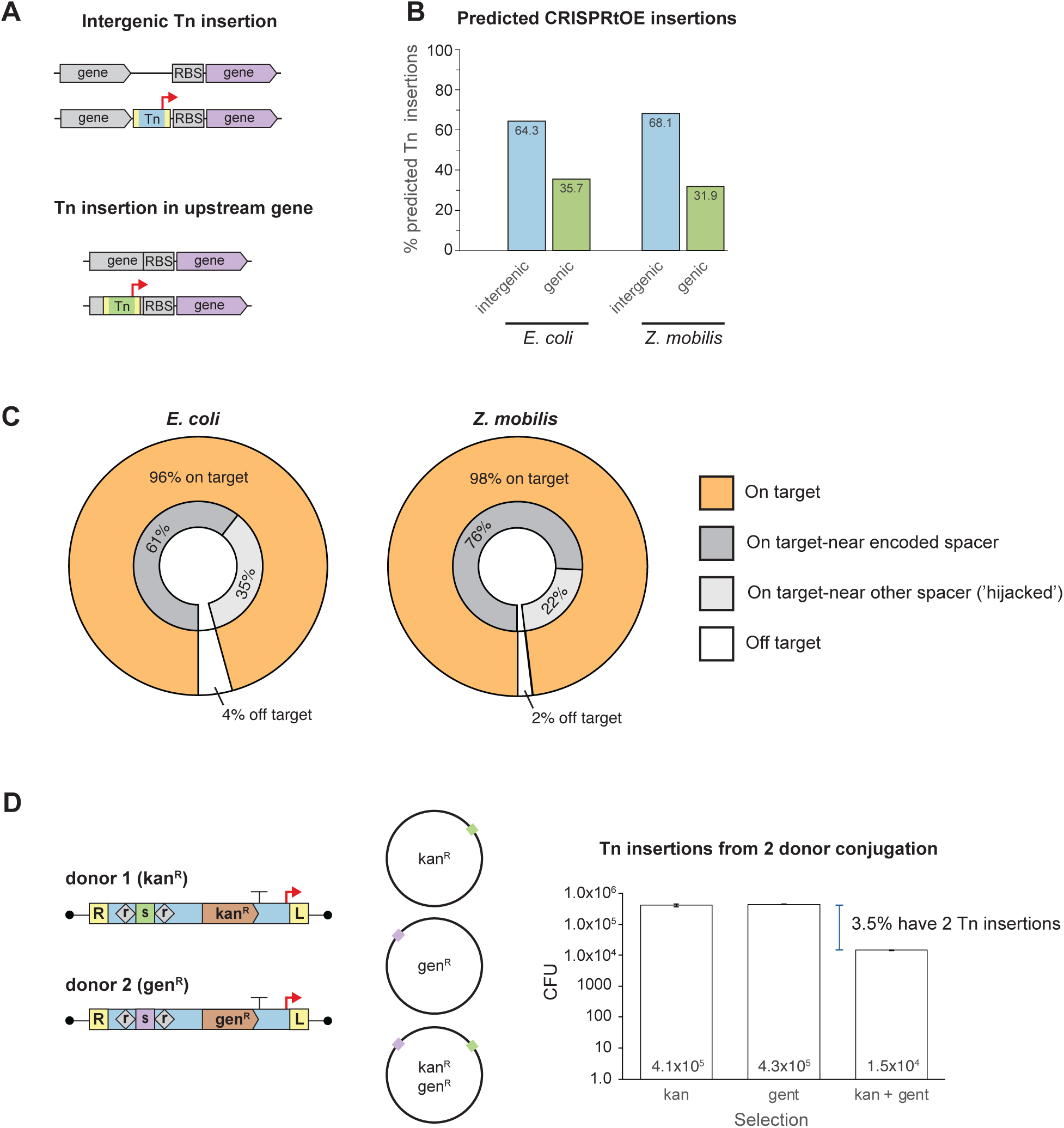
Tn insertion in *E.coli* and *Z. mobilis* CRISPRtOE genome-scale libraries. (A) Schematic of two possible transposon insertion positions: intergenic/entirely between genes or Tn inserted in front of a gene disrupts the end of the gene upstream of it. (B) Percentage of insertions in the *E. coli* and *Z. mobilis* genome-scale CRISPRtOE libraries that are intergenic versus in an upstream gene. (C) Percentage of Tn insertions that are on-(orange) or off- (white) target and percentage on on-target Tn insertions whose position corresponds to the encoded spacer (medium gray) or is instead near the predicted insertion position based on another ’hijacked’ spacer (light gray). (D) Multiplex CRISPRtOE experiment to estimate number of strains that may have multiple Tn insertions. Quad-parental mating was performed with two CRISPRtOE transposon donors with different antibiotic resistance genes (kan^R^ or gen^R^) that were previously shown to effectively target insertion upstream of *folA* or *murA* in *E. coli.* Transconjugants selected on kan, gen, or kan and gen were quantified in three technical replicates and the difference between the number of transconjugants with a single antibiotic resistance phenotype vs. a dual antibiotic resistance phenotype was calculated.

**Figure S11.**
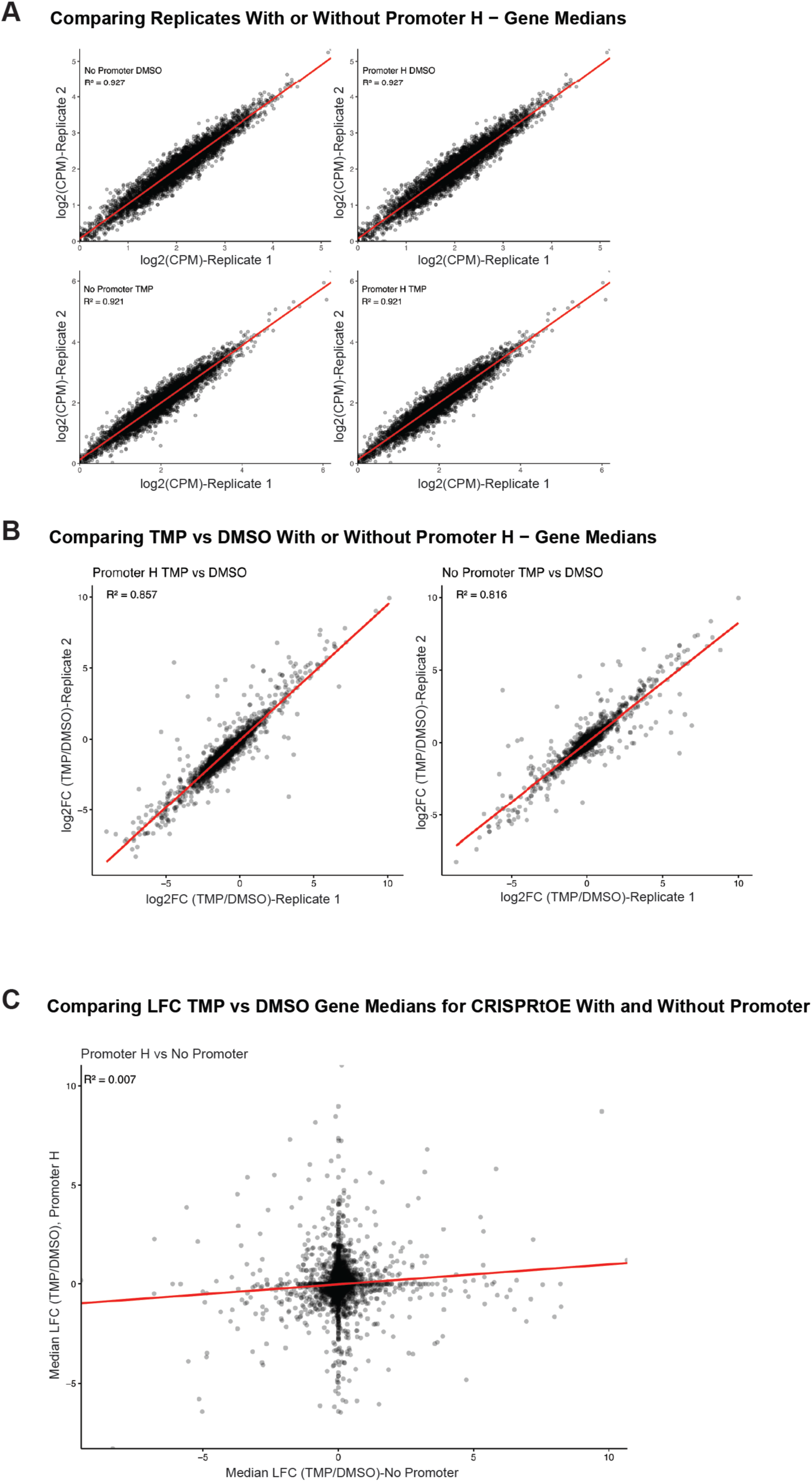
Genome-scale CRISPRtOE reproducibility. (A) Comparison of two biological replicates of CRISPRtOE screen data at the gene level (CPM = counts per million). Either CRISPRtOE library with Promoter H or no promoter with either TMP or DMSO control treatment showing correlation (R^2^ = ∼0.9).(B) Comparison of two biological replicates of the log2FC (FC = fold change) of TMP versus DMSO control treatment for either the CRISPRtOE library with Promoter H or no promoter showing correlation (R^2^ = ∼0.8). (C) Comparison of the median LFC (Log Fold Change) of TMP versus DMSO control treatment for the CRISPRtOE Promoter H versus no promoter libraries showing no correlation (R^2^ = ∼0.007).

**Figure S12.**
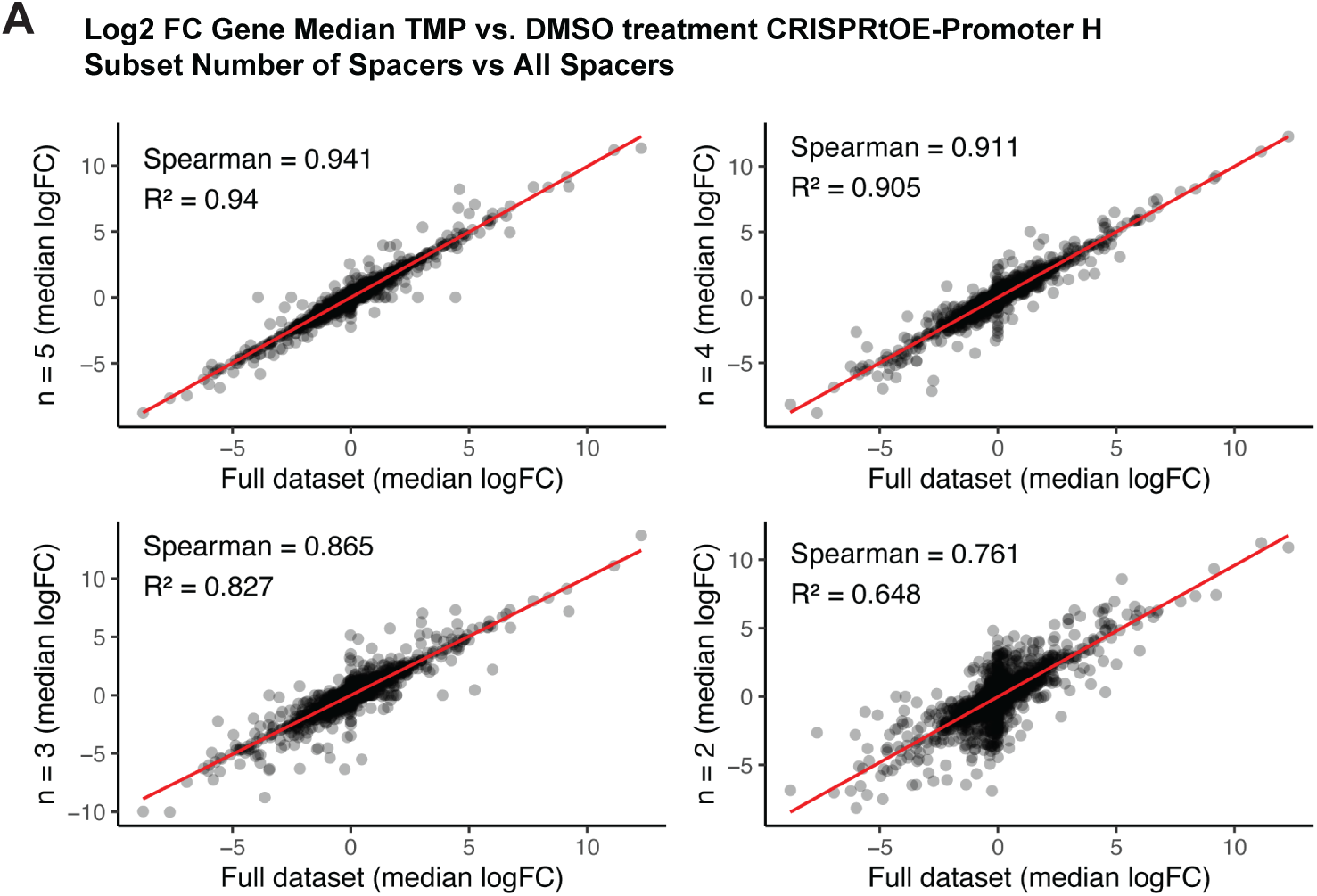
Robustness of CRISPRtOE with fewer analyzed guides. Scatterplots comparing all spacers measured in our study to subsets of 5, 4, 3, or 2 randomly selected spacers per gene.

**Figure S13.**
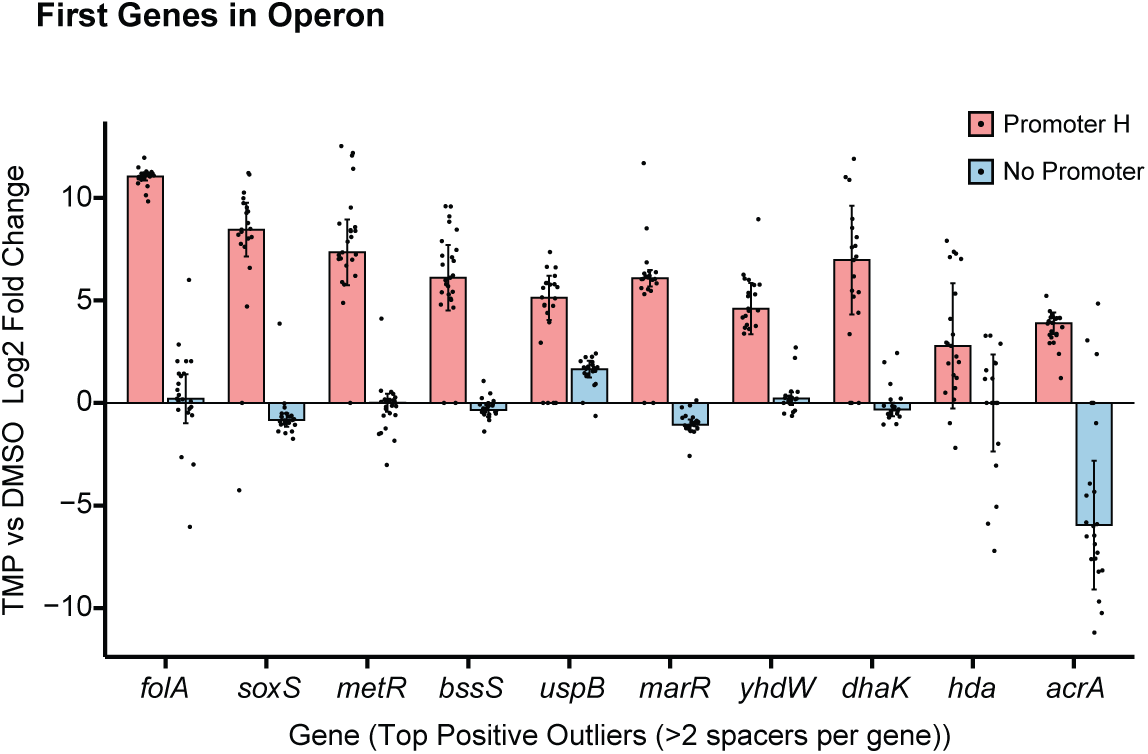
*E. coli* CRISPRtOE TMP screen top positive outlier genes considering only the first gene in the operon. Genes must have at least a 4-fold change in the CRISPRtOE Promoter H data and show a 4-fold difference in a comparison between Promoter H (red) and no promoter (blue) to be considered. Dots represent unique insertions. Error bars represent standard deviation (SD).

**Figure S14.**
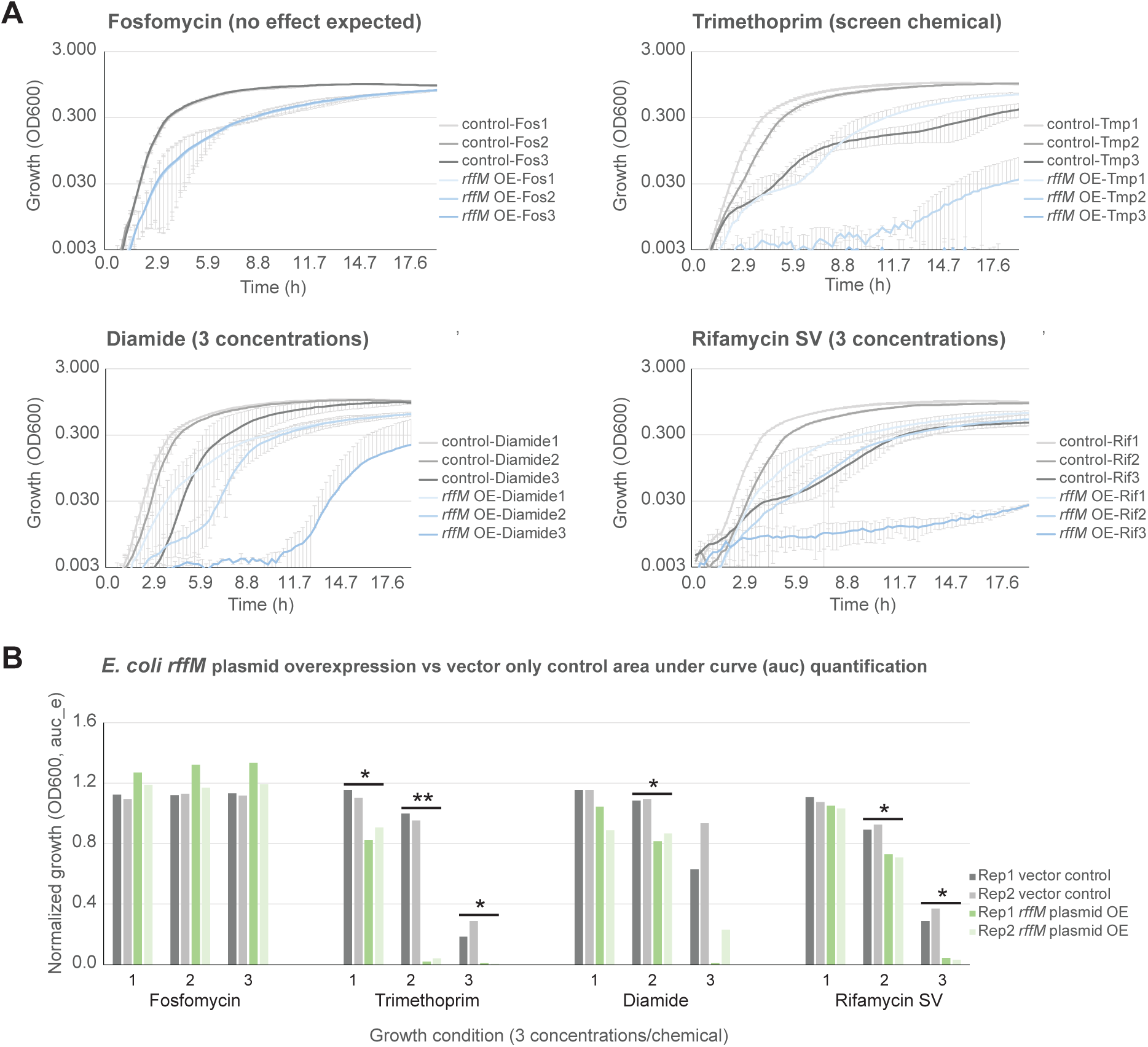
Growth assay of strains overexpressing *rffM* from a plasmid in additional chemical conditions using a Biolog phenotypic array plate PM16 comprised of 24 chemicals with 4 concentrations each. (A) Growth curves of are shown for CRISPRtOE screen chemicals FOS (no response expected) and TMP (response expected), and of two additional chemicals that had a significant response (diamide and rifamycin) for 3 concentrations. Error bars represent SD of duplicate assays. (B) Quantification of area under the curve (auc_e) of growth assays shown in panel (A) using GrowthCurver. Duplicate assays are shown (Rep1 and Rep2). Three concentrations are shown. Significance was determined by a t test (≤0.05(*), ≤0.01(**). Biolog PM16 compounds tested: cefotaxime, phosphomycin, 5-chloro-7-iodo-8-hydroxyquinoline, norfloxacin, sulfanilamide, trimethoprim, dichlofluanid, protamine sulphate, cetylpyridinium chloride, 1-chloro-2,4-dinitrobenzene, diamide, cinoxacin, streptomycin, 5-azacytidine, rifamycin SV, potassium tellurite, sodium selenite, aluminum sulphate, chromium chloride, ferric chloride, L-glutamic-g-hydroxamate, glycine hydroxamate, chloroxylenol, sorbic acid.

**Figure S15.**
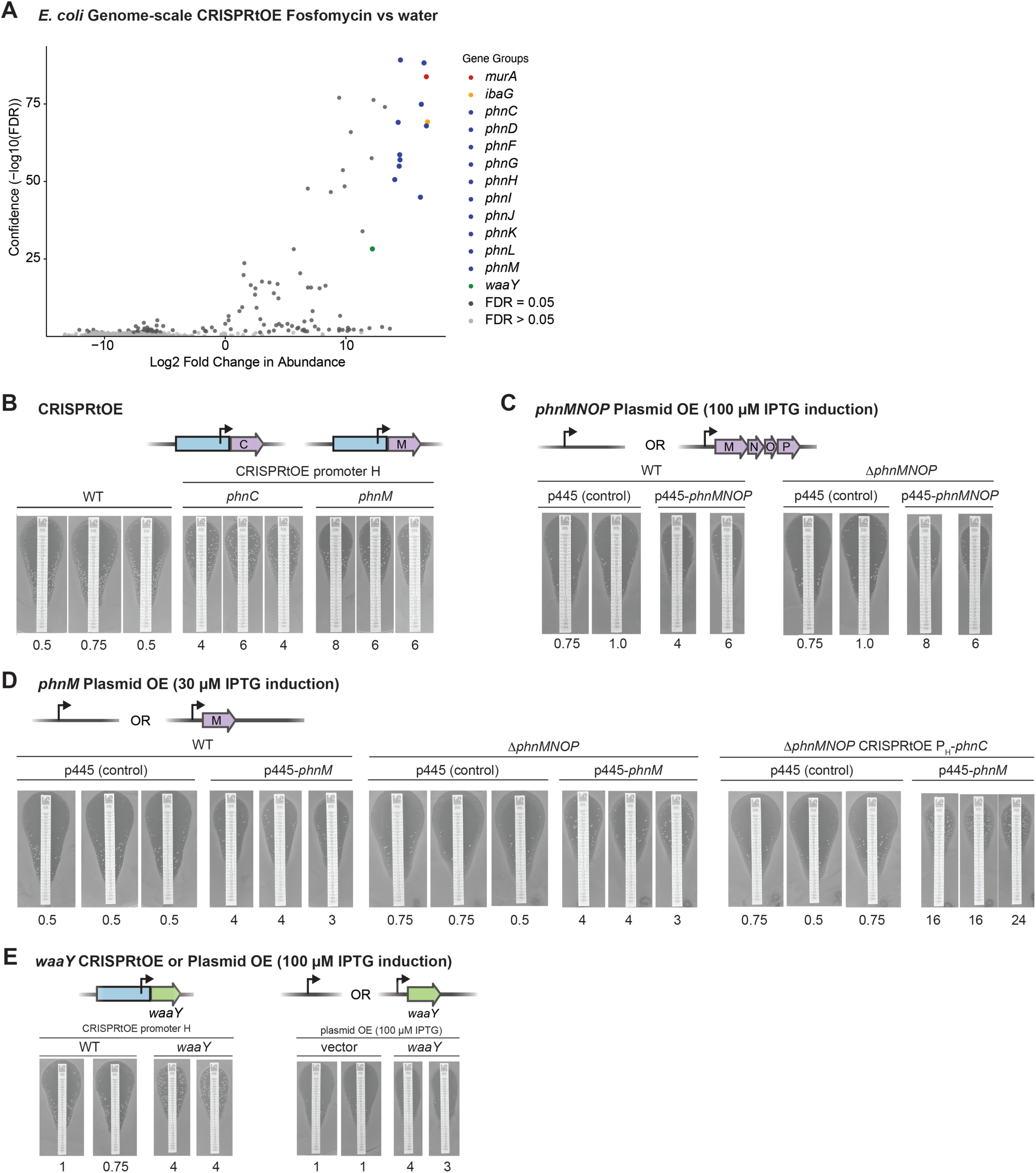
*E. coli* genome scale CRISPRtOE screen with fosfomycin. (A) Volcano plot of gRNA spacer counts in fosfomycin (FOS) versus water control. Each dot represents a strain with a spacer targeting upstream of a gene. Those that are not statistically significant (false discovery rate (FDR) > 0.05) are shown in light grey and significant hits (FDR ≤ 0.05) are shown in dark gray. Several outliers are colored: *murA* (red), *ibaG* (yellow), *phnC-M* (blue), *waaY* (green). (B) MIC test strip (Liofilchem) assay (3 technical replicates) showing resistance to FOS of individual CRISPRtOE-Promoter H strains overexpressing *phnC* or *phnH* compared to WT. (C) MIC test strip (Liofilchem) assay (3 technical replicates) showing resistance to FOS of strains overexpressing *phnMNOP* from a plasmid with an IPTG-inducible promoter in WT or a *Δ..phnMNOP* deletion strain. (D) MIC test strip (Liofilchem) assay (3 technical replicates) showing resistance to FOS of strains overexpressing *phnM* from a plasmid with an IPTG-inducible promoter in WT, *ΔphnMNOP* deletion, or *ΔphnMNOP* deletion plus CRISPRtOE-Promoter H-*phnC* overexpression strain. (E) MIC test strip (Liofilchem) assay (2 technical replicates) showing resistance to FOS of strains overexpressing *waaY* using CRISPRtOE-Promoter H (left) or from a plasmid with an IPTG-inducible promoter (right).

**Figure S16.**
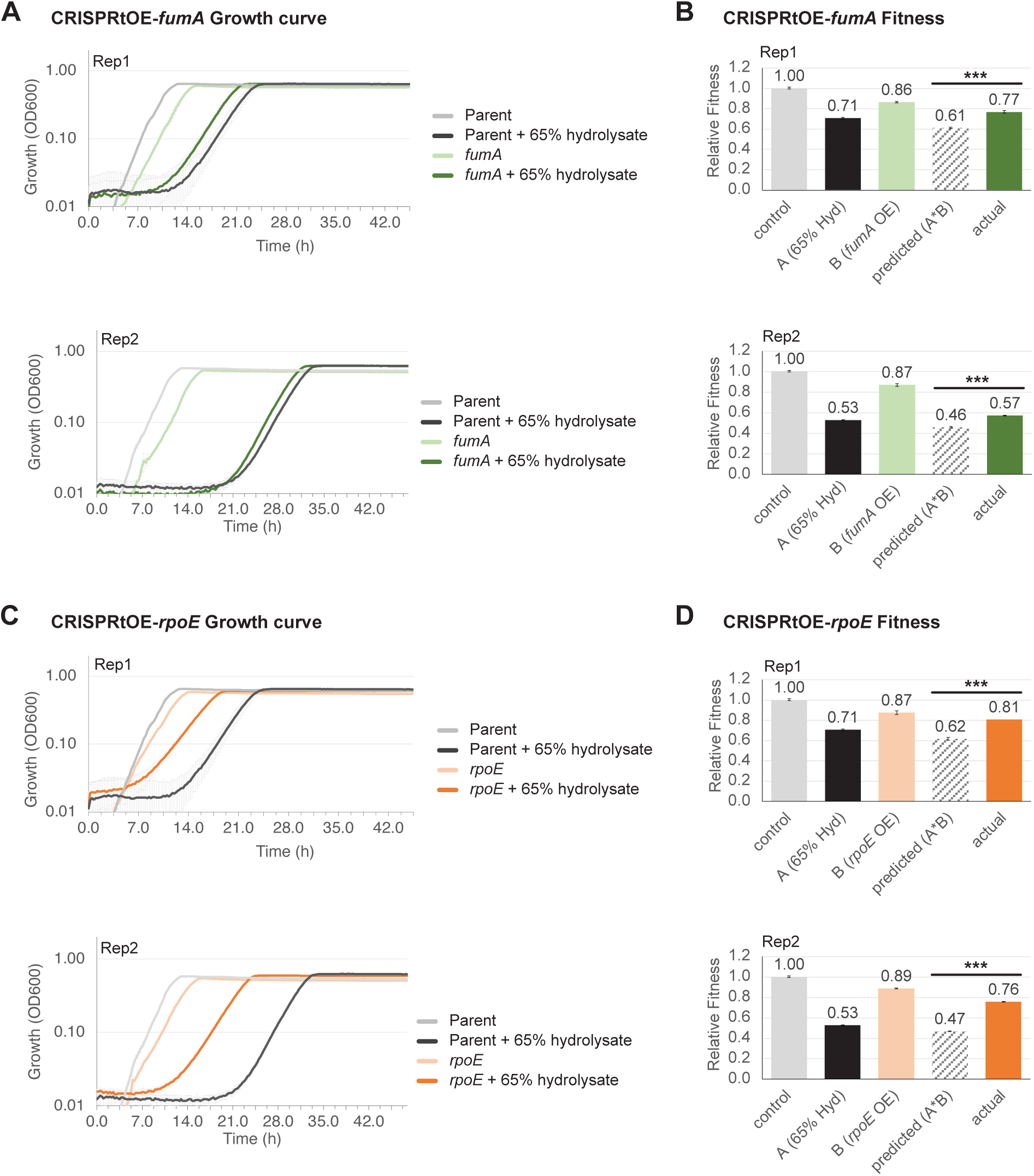
Fitness of *Z*. *mobilis* CRISPRtOE-Promoter H *fumA* and *rpoE* strains. (A) Growth assay of individually constructed CRISPRtOE-Promoter H *fumA* strains versus the parent strain in 65% hydrolysate or ZRDM control. Error bars represent standard deviation (SD) of three independant cultures. (B) Relative fitness of strains shown in panel (A). Growth was quantified by calculating the area under the curve (auc_e) using GrowthCurver. Fitness was normalized to control condition (parent strain in ZRDM). Significance was determined by a t test (≤0.005(***). (C) Growth assay of individually constructed CRISPRtOE-Promoter H *rpoE* strains versus the parent strain in 65% hydrolysate or ZRDM control. Error bars represent standard deviation (SD) of three independant cultures. (D) Relative fitness of strains shown in panel (C). Growth was quantified by calculating the area under the curve (auc_e) using GrowthCurver. Fitness was normalized to control condition (parent strain in ZRDM). Significance was determined by a t test (≤0.005(***).

**Figure S17.**
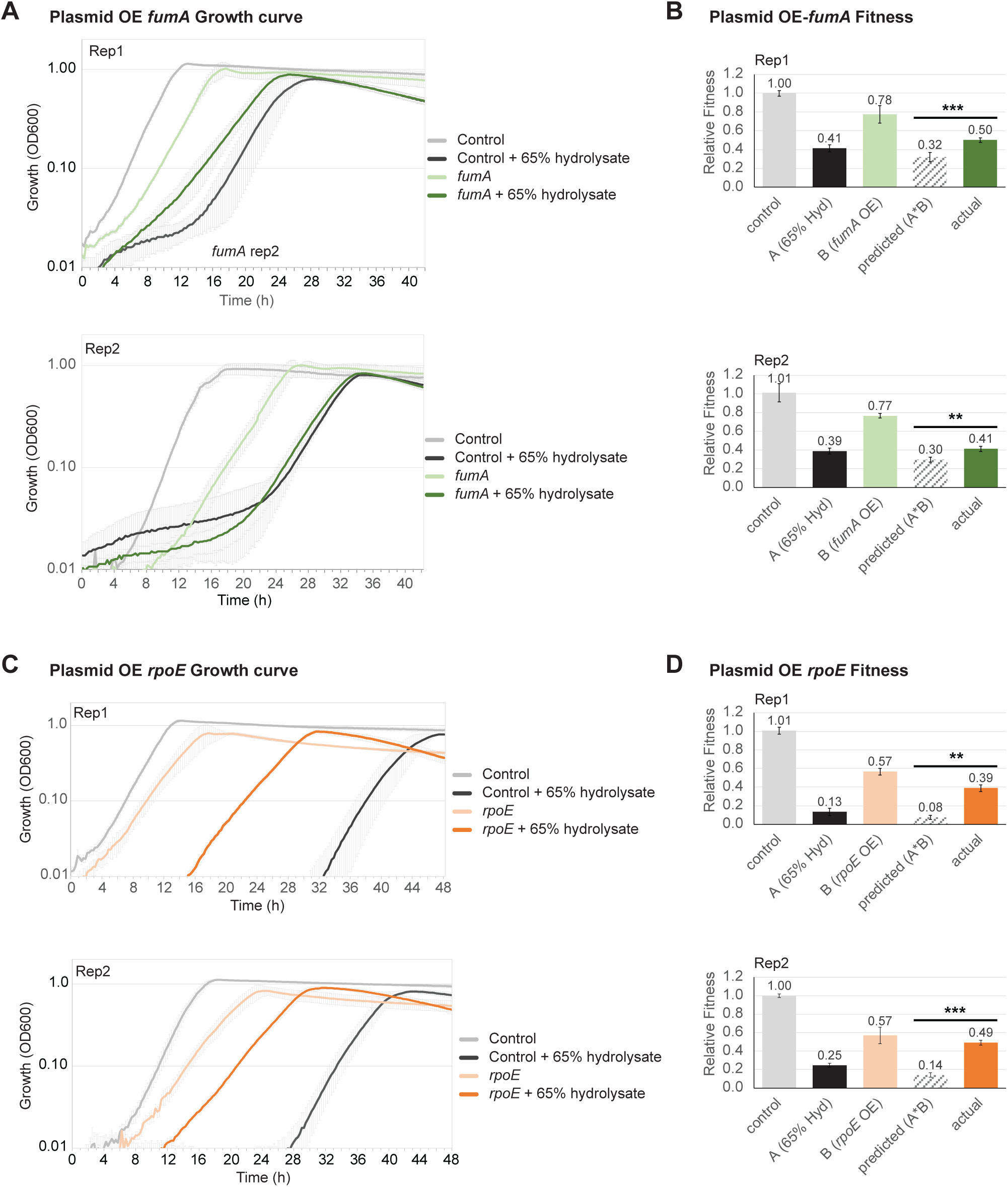
Fitness of *Z*. *mobilis* strains overexpressing *fumA* and *rpoE* from a plasmid. (A) Growth assay of *fumA* plasmid overexpression strains versus a strain with an empty plasmid in 65% hydrolysate or ZRDM control. Error bars represent standard deviation (SD) of three independant cultures. (B) Relative fitness of strains shown in panel (A). Growth was quantified by calculating the area under the curve (auc_e) using GrowthCurver. Fitness was normalized to control condition (parent strain in ZRDM). Significance was determined by a t test (<0.01(**), ≤0.005(***). (C) Growth assay of *rpoE* plasmid overexpression strains versus a strain with an empty plasmid in 65% hydrolysate or ZRDM control. Error bars represent standard deviation (SD) of three independant cultures. (D) Relative fitness of strains shown in panel (C). Growth was quantified by calculating the area under the curve (auc_e) using GrowthCurver. Fitness was normalized to control condition (parent strain in ZRDM). Significance was determined by a t test (<0.01(**), ≤0.005(***).

## References

1. Jost M, Chen Y, Gilbert LA, Horlbeck MA, Krenning L, Menchon G, Rai A, Cho MY, Stern JJ, Prota AE, Kampmann M, Akhmanova A, Steinmetz MO, Tanenbaum ME, Weissman JS. 2017. Combined CRISPRi/a-Based Chemical Genetic Screens Reveal that Rigosertib Is a Microtubule-Destabilizing Agent. Molecular Cell 68:210–223.e6.

2. Kampmann M. 2017. Elucidating drug targets and mechanisms of action by genetic screens in mammalian cells. Chem Commun (Camb) 53:7162–7167.

3. Kampmann M. 2018. CRISPRi and CRISPRa Screens in Mammalian Cells for Precision Biology and Medicine. ACS Chem Biol 13:406–416.

4. Cámara E, Lenitz I, Nygård Y. 2020. A CRISPR activation and interference toolkit for industrial Saccharomyces cerevisiae strain KE6-12. Sci Rep 10:14605.

5. Dominguez AA, Lim WA, Qi LS. 2016. Beyond editing: repurposing CRISPR-Cas9 for precision genome regulation and interrogation. Nat Rev Mol Cell Biol 17:5–15.

6. Jost M, Weissman JS. 2018. CRISPR Approaches to Small Molecule Target Identification. ACS Chem Biol 13:366–375.

7. Lian J, Schultz C, Cao M, HamediRad M, Zhao H. 2019. Multi-functional genome-wide CRISPR system for high throughput genotype-phenotype mapping. Nat Commun 10:5794.

8. Santiago M, Matano LM, Moussa SH, Gilmore MS, Walker S, Meredith TC. 2015. A new platform for ultra-high density Staphylococcus aureus transposon libraries. BMC Genomics 16:252.

9. Kitagawa M, Ara T, Arifuzzaman M, Ioka-Nakamichi T, Inamoto E, Toyonaga H, Mori H. 2005. Complete set of ORF clones of Escherichia coli ASKA library (a complete set of E. coli K-12 ORF archive): unique resources for biological research. DNA Res 12:291–299.

10. Tal S, Paulsson J. 2012. Evaluating quantitative methods for measuring plasmid copy numbers in single cells. Plasmid 67:167–173.

11. Freed EF, Winkler JD, Weiss SJ, Garst AD, Mutalik VK, Arkin AP, Knight R, Gill RT. 2015. Genome-Wide Tuning of Protein Expression Levels to Rapidly Engineer Microbial Traits. ACS Synth Biol 4:1244–1253.

12. Warner JR, Reeder PJ, Karimpour-Fard A, Woodruff LBA, Gill RT. 2010. Rapid profiling of a microbial genome using mixtures of barcoded oligonucleotides. 8. Nat Biotechnol 28:856–862.

13. Mutalik VK, Novichkov PS, Price MN, Owens TK, Callaghan M, Carim S, Deutschbauer AM, Arkin AP. 2019. Dual-barcoded shotgun expression library sequencing for high-throughput characterization of functional traits in bacteria. Nat Commun 10:308.

14. Huang YY, Price MN, Hung A, Gal-Oz O, Ho D, Carion H, Deutschbauer AM, Arkin AP. 2023. Barcoded overexpression screens in gut Bacteroidales identify genes with new roles in carbon utilization and stress resistance. bioRxiv 10.1101/2022.10.10.511384.

15. Mali P, Aach J, Stranges PB, Esvelt KM, Moosburner M, Kosuri S, Yang L, Church GM. 2013. CAS9 transcriptional activators for target specificity screening and paired nickases for cooperative genome engineering. Nat Biotechnol 31:833–838.

16. Lee DJ, Minchin SD, Busby SJW. 2012. Activating transcription in bacteria. Annu Rev Microbiol 66:125–152.

17. Bikard D, Jiang W, Samai P, Hochschild A, Zhang F, Marraffini LA. 2013. Programmable repression and activation of bacterial gene expression using an engineered CRISPR-Cas system. Nucl Acids Res gkt520.

18. Kiattisewee C, Karanjia AV, Legut M, Daniloski Z, Koplik SE, Nelson J, Kleinstiver BP, Sanjana NE, Carothers JM, Zalatan JG. 2022. Expanding the Scope of Bacterial CRISPR Activation with PAM-Flexible dCas9 Variants. ACS Synth Biol 11:4103–4112.

19. Ho H-I, Fang JR, Cheung J, Wang HH. 2020. Programmable CRISPR-Cas transcriptional activation in bacteria. Mol Syst Biol 16:e9427.

20. Peters JE, Makarova KS, Shmakov S, Koonin EV. 2017. Recruitment of CRISPR-Cas systems by Tn7-like transposons. PNAS 114:E7358–E7366.

21. Vo PLH, Ronda C, Klompe SE, Chen EE, Acree C, Wang HH, Sternberg SH. 2021. CRISPR RNA-guided integrases for high-efficiency, multiplexed bacterial genome engineering. Nat Biotechnol 39:480–489.

22. Rubin BE, Diamond S, Cress BF, Crits-Christoph A, Lou YC, Borges AL, Shivram H, He C, Xu M, Zhou Z, Smith SJ, Rovinsky R, Smock DCJ, Tang K, Owens TK, Krishnappa N, Sachdeva R, Barrangou R, Deutschbauer AM, Banfield JF, Doudna JA. 2022. Species- and site-specific genome editing in complex bacterial communities. Nat Microbiol 7:34–47.

23. Chen W, Ren Z-H, Tang N, Chai G, Zhang H, Zhang Y, Ma J, Wu Z, Shen X, Huang X, Luo G-Z, Ji Q. 2021. Targeted genetic screening in bacteria with a Cas12k-guided transposase. Cell Rep 36:109635.

24. Trujillo Rodríguez L, Ellington AJ, Reisch CR, Chevrette MG. 2023. CRISPR-Associated Transposase for Targeted Mutagenesis in Diverse Proteobacteria. ACS Synth Biol 12:1989–2003.

25. Roberts A, Nethery MA, Barrangou R. 2022. Functional characterization of diverse type I-F CRISPR-associated transposons. Nucleic Acids Res 50:11670–11681.

26. Hsieh S-C, Peters JE. 2023. Discovery and characterization of novel type I-D CRISPR-guided transposons identified among diverse Tn7-like elements in cyanobacteria. Nucleic Acids Res 51:765–782.

27. Walker MWG, Klompe SE, Zhang DJ, Sternberg SH. 2023. Novel molecular requirements for CRISPR RNA-guided transposition. Nucleic Acids Res 51:4519–4535.

28. Gelsinger DR, Vo PLH, Klompe SE, Ronda C, Wang H, Sternberg SH. 2023. Bacterial genome engineering using CRISPR RNA-guided transposases. bioRxiv 2023.03.18.533263.

29. Hoffmann FT, Kim M, Beh LY, Wang J, Vo PLH, Gelsinger DR, George JT, Acree C, Mohabir JT, Fernández IS, Sternberg SH. 2022. Selective TnsC recruitment enhances the fidelity of RNA-guided transposition. Nature 609:384–393.

30. Halpin-Healy TS, Klompe SE, Sternberg SH, Fernández IS. 2020. Structural basis of DNA targeting by a transposon-encoded CRISPR-Cas system. Nature 577:271–274.

31. Klompe SE, Vo PLH, Halpin-Healy TS, Sternberg SH. 2019. Transposon-encoded CRISPR–Cas systems direct RNA-guided DNA integration. Nature 571:219.

32. Garza Elizondo AM, Chappell J. 2024. Targeted Transcriptional Activation Using a CRISPR-Associated Transposon System. ACS Synth Biol 13:328–336.

33. Cain AK, Barquist L, Goodman AL, Paulsen IT, Parkhill J, van Opijnen T. 2020. A decade of advances in transposon-insertion sequencing. Nat Rev Genet 21:526–540.

34. Petassi MT, Hsieh S-C, Peters JE. 2020. Guide RNA Categorization Enables Target Site Choice in Tn7-CRISPR-Cas Transposons. Cell 183:1757–1771.e18.

35. Banta AB, Enright AL, Siletti C, Peters JM. 2020. A High-efficacy CRISPRi System for Gene Function Discovery in Zymomonas mobilis. Appl Environ Microbiol 10.1128/AEM.01621-20.

36. Hall AN, Hall BW, Kinney KJ, Olsen GG, Banta AB, Noguera DR, Donohue TJ, Peters JM. 2023. Tools for Genetic Engineering and Gene Expression Control in Novosphingobium aromaticivorans and Rhodobacter sphaeroides. bioRxiv 2023.08.25.554875.

37. Enright AL, Banta AB, Ward RD, Rivera Vazquez J, Felczak MM, Wolfe MB, TerAvest MA, Amador-Noguez D, Peters JM. 2023. The genetics of aerotolerant growth in an alphaproteobacterium with a naturally reduced genome. mBio e0148723.

38. Rice LB. 2008. Federal funding for the study of antimicrobial resistance in nosocomial pathogens: no ESKAPE. J Infect Dis 197:1079–1081.

39. Fenster JA, Werner AZ, Tay JW, Gillen M, Schirokauer L, Hill NC, Watson A, Ramirez KJ, Johnson CW, Beckham GT, Cameron JC, Eckert CA. 2022. Dynamic and single cell characterization of a CRISPR-interference toolset in Pseudomonas putida KT2440 for β-ketoadipate production from p-coumarate. Metab Eng Commun 15:e00204.

40. Hau HH, Gralnick JA. 2007. Ecology and biotechnology of the genus Shewanella. Annu Rev Microbiol 61:237–258.

41. Prasad NK, Seiple IB, Cirz RT, Rosenberg OS. 2022. Leaks in the Pipeline: a Failure Analysis of Gram-Negative Antibiotic Development from 2010 to 2020. Antimicrobial Agents and Chemotherapy 66:e00054–22.

42. Palmer AC, Kishony R. 2014. Opposing effects of target overexpression reveal drug mechanisms. Nat Commun 5:4296.

43. Couce A, Briales A, Rodríguez-Rojas A, Costas C, Pascual A, Blázquez J. 2012. Genomewide overexpression screen for fosfomycin resistance in Escherichia coli: MurA confers clinical resistance at low fitness cost. Antimicrob Agents Chemother 56:2767– 2769.

44. Ason B, Reznikoff WS. 2004. DNA sequence bias during Tn5 transposition. J Mol Biol 335:1213–1225.

45. Lee J-H, Lee K-L, Yeo W-S, Park S-J, Roe J-H. 2009. SoxRS-mediated lipopolysaccharide modification enhances resistance against multiple drugs in Escherichia coli. J Bacteriol 191:4441–4450.

46. Seo SW, Kim D, Szubin R, Palsson BO. 2015. Genome-wide Reconstruction of OxyR and SoxRS Transcriptional Regulatory Networks under Oxidative Stress in Escherichia coli K-12 MG1655. Cell Rep 12:1289–1299.

47. Jones KM, Guest JR, Woods DD. 1961. Folic acid and the synthesis of methionine by extracts of Escherichia coli. Biochem J 79:566–574.

48. Weissbach H, Brot N. 1991. Regulation of methionine synthesis in Escherichia coli. Mol Microbiol 5:1593–1597.

49. Saint-Girons I, Duchange N, Zakin MM, Park I, Margarita D, Ferrara P, Cohen GN. 1983. Nucleotide sequence of metF, the E. coli structural gene for 5-10 methylene tetrahydrofolate reductase and of its control region. Nucleic Acids Res 11:6723–6732.

50. Girgis HS, Hottes AK, Tavazoie S. 2009. Genetic architecture of intrinsic antibiotic susceptibility. PLoS One 4:e5629.

51. Nichols RJ, Sen S, Choo YJ, Beltrao P, Zietek M, Chaba R, Lee S, Kazmierczak KM, Lee KJ, Wong A, Shales M, Lovett S, Winkler ME, Krogan NJ, Typas A, Gross CA. 2011. Phenotypic landscape of a bacterial cell. Cell 144:143–156.

52. Giladi M, Altman-Price N, Levin I, Levy L, Mevarech M. 2003. FolM, a new chromosomally encoded dihydrofolate reductase in Escherichia coli. J Bacteriol 185:7015–7018.

53. Rao TVP, Kuzminov A. 2020. Exopolysaccharide defects cause hyper-thymineless death in Escherichia coli via massive loss of chromosomal DNA and cell lysis. Proc Natl Acad Sci U S A 117:33549–33560.

54. Metcalf WW, Wanner BL. 1991. Involvement of the Escherichia coli phn (psiD) gene cluster in assimilation of phosphorus in the form of phosphonates, phosphite, Pi esters, and Pi. J Bacteriol 173:587–600.

55. Turner AK, Yasir M, Bastkowski S, Telatin A, Page AJ, Charles IG, Webber MA. 2020. A genome-wide analysis of Escherichia coli responses to fosfomycin using TraDIS-Xpress reveals novel roles for phosphonate degradation and phosphate transport systems. J Antimicrob Chemother 75:3144–3151.

56. Yethon JA, Heinrichs DE, Monteiro MA, Perry MB, Whitfield C. 1998. Involvement of waaY, waaQ, and waaP in the modification of Escherichia coli lipopolysaccharide and their role in the formation of a stable outer membrane. J Biol Chem 273:26310–26316.

57. Jacobson TB, Adamczyk PA, Stevenson DM, Regner M, Ralph J, Reed JL, Amador-Noguez D. 2019. 2H and 13C metabolic flux analysis elucidates in vivo thermodynamics of the ED pathway in Zymomonas mobilis. Metabolic Engineering 54:301–316.

58. Erickson JW, Gross CA. 1989. Identification of the sigma E subunit of Escherichia coli RNA polymerase: a second alternate sigma factor involved in high-temperature gene expression. Genes Dev 3:1462–1471.

59. Newman JD, Falkowski MJ, Schilke BA, Anthony LC, Donohue TJ. 1999. The Rhodobacter sphaeroides ECF sigma factor, sigma(E), and the target promoters cycA P3 and rpoE P1. J Mol Biol 294:307–320.

60. Partridge SR, Kwong SM, Firth N, Jensen SO. 2018. Mobile Genetic Elements Associated with Antimicrobial Resistance. Clin Microbiol Rev 31:e00088–17.

61. Martínez-García E, Aparicio T, de Lorenzo V, Nikel PI. 2014. New transposon tools tailored for metabolic engineering of gram-negative microbial cell factories. Front Bioeng Biotechnol 2:46.

62. Yang Q, Yang Y, Tang Y, Wang X, Chen Y, Shen W, Zhan Y, Gao J, Wu B, He M, Chen S, Yang S. 2020. Development and characterization of acidic-pH-tolerant mutants of Zymomonas mobilis through adaptation and next-generation sequencing-based genome resequencing and RNA-Seq. Biotechnol Biofuels 13:144.

63. Dubois L, Vettiger A, Buss JA, Bernhardt TG. 2025. Using fluorescently labeled wheat germ agglutinin to track lipopolysaccharide transport to the outer membrane in Escherichia coli. mBio 16:e0395024.

64. Kamat SS, Williams HJ, Raushel FM. 2011. Intermediates in the transformation of phosphonates to phosphate by bacteria. Nature 480:570–573.

65. Amstrup SK, Ong SC, Sofos N, Karlsen JL, Skjerning RB, Boesen T, Enghild JJ, Hove-Jensen B, Brodersen DE. 2023. Structural remodelling of the carbon-phosphorus lyase machinery by a dual ABC ATPase. Nat Commun 14:1001.

66. Tenaillon O, Rodríguez-Verdugo A, Gaut RL, McDonald P, Bennett AF, Long AD, Gaut BS. 2012. The molecular diversity of adaptive convergence. Science 335:457–461.

67. Banta AB, Ward RD, Tran JS, Bacon EE, Peters JM. 2020. Programmable Gene Knockdown in Diverse Bacteria Using Mobile-CRISPRi. Curr Protoc Microbiol 59:e130.

68. Sharan SK, Thomason LC, Kuznetsov SG, Court DL. 2009. Recombineering: a homologous recombination-based method of genetic engineering. Nat Protoc 4:206–223.

69. Thomason LC, Costantino N, Court DL. 2007. E. coli genome manipulation by P1 transduction. Curr Protoc Mol Biol Chapter 1:1.17.1–1.17.8.

70. Cherepanov PP, Wackernagel W. 1995. Gene disruption in Escherichia coli: TcR and KmR cassettes with the option of Flp-catalyzed excision of the antibiotic-resistance determinant. Gene 158:9–14.

71. Peters JM, Koo B-M, Patino R, Heussler GE, Hearne CC, Qu J, Inclan YF, Hawkins JS, Lu CHS, Silvis MR, Harden MM, Osadnik H, Peters JE, Engel JN, Dutton RJ, Grossman AD, Gross CA, Rosenberg OS. 2019. Enabling genetic analysis of diverse bacteria with Mobile-CRISPRi. Nature Microbiology 4:244–250.

72. Livak KJ, Schmittgen TD. 2001. Analysis of relative gene expression data using real-time quantitative PCR and the 2(-Delta Delta C(T)) Method. Methods 25:402–408.

73. Martin M. 2011. Cutadapt removes adapter sequences from high-throughput sequencing reads. 1. EMBnet.journal 17:10–12.

74. Langmead B, Trapnell C, Pop M, Salzberg SL. 2009. Ultrafast and memory-efficient alignment of short DNA sequences to the human genome. Genome Biology 10:R25.

75. Li H, Handsaker B, Wysoker A, Fennell T, Ruan J, Homer N, Marth G, Abecasis G, Durbin R, 1000 Genome Project Data Processing Subgroup. 2009. The Sequence Alignment/Map format and SAMtools. Bioinformatics 25:2078–2079.

76. Bushnell B. 2014. BBMap: A Fast, Accurate, Splice-Aware Aligner.

